# Droplet-based combinatorial indexing for massive scale single-cell epigenomics

**DOI:** 10.1101/612713

**Authors:** Caleb A. Lareau, Fabiana M. Duarte, Jennifer G. Chew, Vinay K. Kartha, Zach D. Burkett, Andrew S. Kohlway, Dmitry Pokholok, Martin J. Aryee, Frank J. Steemers, Ronald Lebofsky, Jason D. Buenrostro

## Abstract

While recent technical advancements have facilitated the mapping of epigenomes at single-cell resolution, the throughput and quality of these methods have limited the widespread adoption of these technologies. Here, we describe a droplet microfluidics platform for single-cell assay for transposase accessible chromatin (scATAC-seq) for high-throughput single-cell profiling of chromatin accessibility. We use this approach for the unbiased discovery of cell types and regulatory elements within the mouse brain. Further, we extend the throughput of this approach by pairing combinatorial indexing with droplet microfluidics, enabling single-cell studies at a massive scale. With this approach, we measure chromatin accessibility across resting and stimulated human bone marrow derived cells to reveal changes in the *cis*- and *trans-* regulatory landscape across cell types and upon stimulation conditions at single-cell resolution. Altogether, we describe a total of 502,207 single-cell profiles, demonstrating the scalability and flexibility of this droplet-based platform.

## Introduction

While the primary sequence of the eukaryotic genome is largely invariant across cells in an organism, the quantitative expression of genes is tightly regulated to define the functional identity of cells. Eukaryotic cells use diverse mechanisms to regulate gene expression, including an immense repertoire (>10^6^) of DNA regulatory elements^1,2^. These DNA regulatory elements are established and maintained by the combinatorial binding of transcription factors (TFs) and chromatin remodelers, which together function to recruit transcriptional machinery and drive cell-type specific gene expression^3,4^. DNA regulatory elements, characterized by their functional roles (promoter, enhancer, insulator, etc.), are marked by a diverse array of histone and DNA modifications^4^. Importantly, both classical observations^5^ and recent genome-wide efforts^2^ have observed that active regulatory elements are canonically nucleosome free and accessible to transcriptional machinery. Thus, methods that measure chromatin accessibility based on sensitivity to enzymatic digestion followed by sequencing^6–8^ provide an integrated map of chromatin states which encompass a diverse repertoire of functional regulatory elements^2,5^.

Methods to assay chromatin accessibility genome-wide have been used for a variety of applications including the discovery of i) cell type-specific *cis*-regulatory elements, ii) master TFs that shape the regulatory landscape, or iii) mechanisms for disease-relevant non-coding genetic variation^2,9,10^. However, these “epigenomic” approaches are generally applied to bulk samples, limiting their resolution into the regulatory diversity underlying heterogeneous cell populations. In parallel, methods to measure the transcriptomes of single-cells have been used to discover new cell types^11^ and new functional cell states^12,13^, and provide additional motivation for the development of tools to measure chromatin states at single-cell resolution^14^.

Technological innovations have enabled the development of single-cell epigenomic methods^14–16^; however, these approaches remain relatively low-throughput and high-cost. Assay for Transposase Accessible chromatin (ATAC-seq)^8,17^is particularly promising for single-cell studies due to the relative simplicity of the experimental protocol, and widespread use. Previous efforts have adapted ATAC-seq to profile chromatin accessibility in single-cells, either by individually isolating cells^18^ or by combinatorial addition of DNA barcodes^19^, to enable *de novo* deconvolution of cell types and the discovery of cell-type specific regulatory factors^20,21^. However, these current methods for single-cell ATAC-seq (scATAC-seq)^18,19^ remain either relatively low-throughput (100s to 1,000s of cells/experiment) or provide low-complexity data (1,000s of fragments per cell). Therefore, new methods for sensitive, scalable, and high-throughput profiling are needed to measure the full repertoire of regulatory diversity across normal and diseased tissues.

To meet the challenges of assaying chromatin states in the breadth and depth of complex cell populations within tissues, we report the development of a droplet-based scATAC-seq assay. In brief, our approach utilizes a droplet microfluidic device to individually isolate and barcode transposed single-cells. We demonstrate that this approach results in significantly higher data quality than existing methods, and describe an approach to improve cell throughput and cell capture efficiency by super-loading barcoded beads into droplets. Further, we extend this droplet barcoding approach by combining this method with barcoded transposition^19^ followed by super-loading cells into droplets, to develop droplet-based single-cell combinatorial indexing for ATAC-seq (dsciATAC-seq), providing chromatin accessibility profiles at significantly improved throughput. As a proof of principle, we apply these approaches to generate accessibility profiles of 502,207 cells, which includes i) a reference map of chromatin accessibility in the mouse brain (38,737 cells), and ii) an unbiased map of human hematopoietic states in the bone marrow (60,495 cells), isolated cell populations from bone marrow and blood (52,873 cells), and bone marrow cells in response to stimulation (75,958 cells). These unbiased chromatin accessibility profiles provide new insights into the regulators defining cells within these tissues. Further, we find that pooled stimulus of human bone marrow derived cells uncovers mechanistic insights driving genetic variants leading to human disease. Overall, this new approach for high-throughput single-cell epigenomics charts a clear course towards obtaining an epigenomic atlas across normal tissues, and provides new opportunities for single-cell epigenomic profiling at a massive scale.

## Results

### scATAC-seq implemented on a droplet microfluidic device

In this work we describe a method for single-cell chromatin accessibility profiling using droplet microfluidics and ATAC-seq. Consistent with previously described methods for bulk ATAC-seq, nuclei are first transposed using Tn5 transposase to integrate sequencing adapters into regions of open chromatin^8,17,22^. Importantly, previous studies have described that transposed nuclei and DNA remain intact following transposition^19,23^. We therefore leverage this finding and use intact transposed nuclei as input material into a droplet microfluidics device, which co-encapsulates transposed chromatin with PCR reagents and barcoded beads into a single droplet (**Fig. 1a**). Each bead contains clonal copies of oligonucleotides that encode a common PCR primer sequence and a bead-specific DNA barcode. Following co-encapsulation, we perform droplet PCR to add cell-identifying DNA barcodes to transposed chromatin, and the resulting pool of PCR products are then collected and prepared for sequencing. To further facilitate a robust and high-throughput platform, we optimized Tn5 transposase and droplet PCR conditions to improve the total number of nuclear genomic fragments and reduce the relative frequency of mitochondrial reads in sample preparation (**Fig. S1**). We refer to this droplet-based scATAC-seq platform as dscATAC-seq.

**Figure 1.**
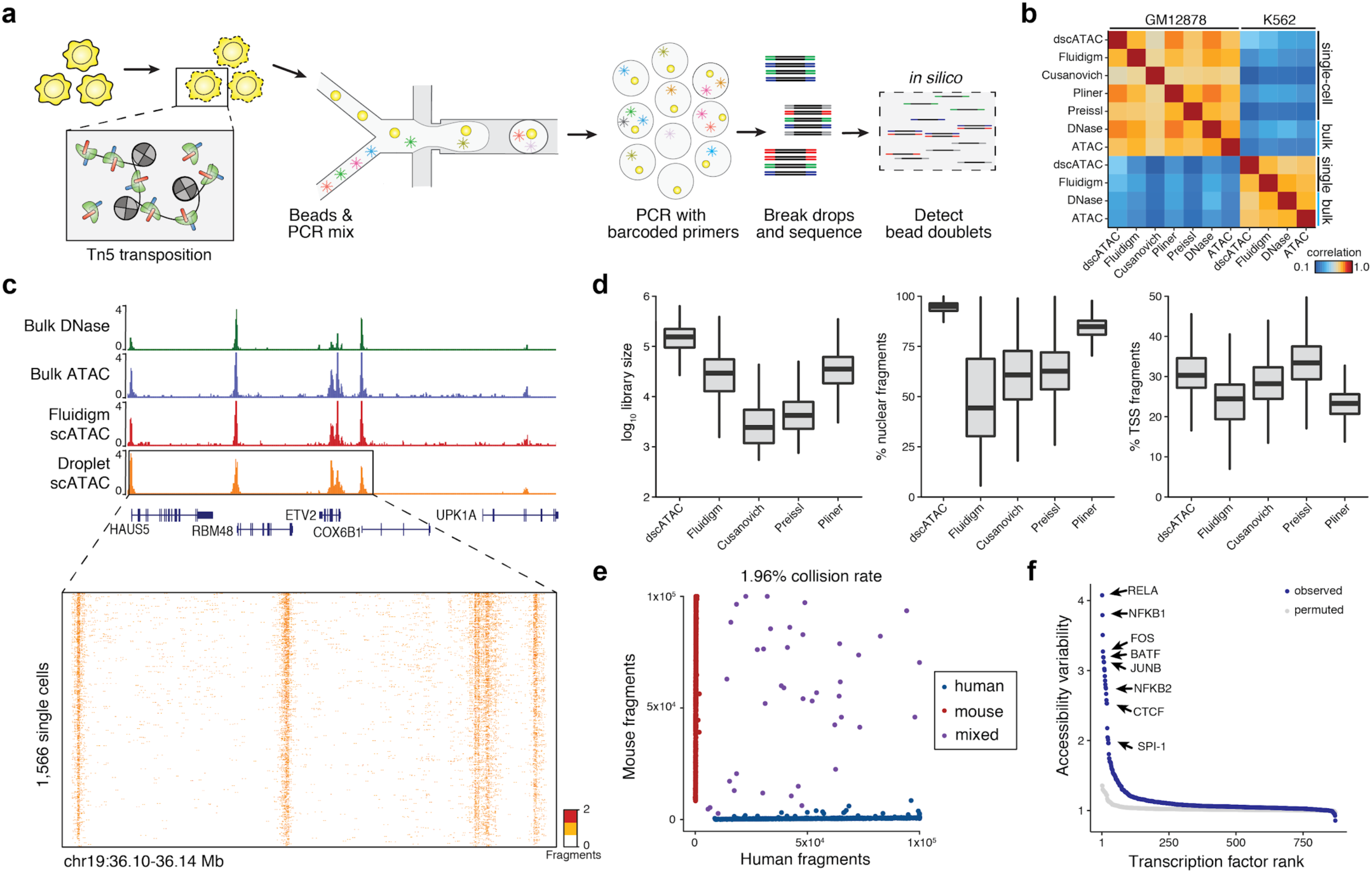
dscATAC-seq enables high-resolution characterization of open chromatin regions in single cells. (**a**) Schematic of technology. Cells are transposed with Tn5 transposase, transposed chromatin is then barcoded and amplified in a microfluidic device. (**b**) Spearman correlation of reads in chromatin accessibility peaks across bulk and single-cell technologies for GM12878 and K562 cells. (**c**) Comparison of the aggregate chromatin accessibility profiles from GM12878 cells using different technologies, and visualization of single-cell chromatin accessibility profiles from dscATAC-seq. (**d**) Quality metrics of scATAC-seq methods for GM12878 cells. Center line, median; box limits, first and third quartiles; whiskers, 1.5x interquartile range. Median library size for dscATAC-seq was 165,204 reads (left panel, all reads reported are passing quality filters), compared to profiles generated from the Fluidigm C1^18^ (50,443 reads) and a recently optimized sciATAC-seq method (46,730 reads, “Pliner”^25^). Median fraction of mapped nuclear fragments for dscATAC-seq is 95% (middle panel). (**e**) The number of unique fragments aligning to human or mouse genomes using human (GM12878) and mouse (3T3) cells at 800 beads/μL. (**f**) Rank sorted variability across transcription factor motifs within the GM12878 dscATAC-seq profiles.

To improve cell capture and throughput, we developed a joint experimental and computational strategy to super-load beads into droplets. Our computational strategy, which we call the **b**ead-based sc**A**TAC **p**rocessing (bap), determines barcodes with a high overlap of aligned fragment positions to identify and merge bead barcodes within a common droplet (**Fig. S2a-d**), which enables loading beads at higher density to increase the number of droplets with one or more beads (**Fig. S2e-f**). We verified this computational approach to be highly accurate using a library of random oligonucleotides as true positives (**Fig. S2g**, see Methods). We also found consistent experimental results across a range of bead concentrations without loss of data quality (**Fig. S2h-k**). To compare the efficacy of our approach, we uniformly processed cell line data (GM12878 and K562) generated using dscATAC-seq and from four other recently published approaches^18,19,24,25^. We found that bulk ATAC-seq^8^, DNase-seq^2^ and the aggregate chromatin accessibility across the different single-cell technologies^18,19,24,25^ were highly correlated (**Fig. 1b,c**). However, our dscATAC-seq method achieved improved library complexity per cell, proportion of reads mapping to the nuclear genome, and number of cells per experiment (**Fig. 1d** and **Fig. S3a,b**) – common quality metrics for scATAC-seq experiments. Further, we also observed < 2% collision rate with 5,600 cells as input, yielding 2,040 (800 beads/μL) to 2,593 (5,000 beads/μL) cells passing filter (**Fig. 1e** and **Fig. S2j,k**). We note that this approach for super loading beads improved cell throughput (approximately 300 to 2,500 cells) and cell capture rates (approximately 5% to 45%), and note that our estimated collision rate (< 2%) is considerably lower than other previously described high-throughput sciATAC-seq methods^19,24^(> 5%) (**Fig. S3c**). Using this method, we also find similar variation in TF motif activity across single GM12878 cells as previously reported^18,26^ (**Fig. 1f**). Taken together, our new methodology provides an approach for high-resolution profiling of chromatin accessibility across thousands of single cells.

### Epigenomic diversity of the adult mouse brain

We sought to determine whether our approach could be applied to large-scale efforts to identify cell types within complex tissues *de novo*. Thus, we applied the dscATAC-seq platform to whole brain tissues derived from two mice, collecting an average of 3,874 cells per well, for a total of 38,737 cells passing quality thresholds. Cells passing quality filters had a median of 30,020 unique nuclear reads, 58.7% reads in peaks and an average of 2 bead barcodes per cell (**Fig. S4a**).

To characterize differences in chromatin accessibility across cell types, we first reduced dimensionality of our mouse brain profiles by computing K-mer deviation scores (7-mers) using the chromVAR algorithm^26^. Cell clusters are identified using the Louvain modularity method built from a cell nearest-neighbor graph using the 7-mer scores, which uncovered 25 cell clusters. We then use these 7-mer features to subsequently map each cell to a two-dimensional representation using t-Distributed Stochastic Neighbor Embedding (t-SNE) (**Fig. 2a**). Importantly, these clusters are largely uncorrelated with known technical confounders (**Fig. S4b-e**). For comparison with previous methods, we also analyzed published sciATAC-seq data from two mouse brains^27^, where we identified 13 clusters using this computational approach (**Fig. S4f**), which we attribute to the fewer cells assayed (5,744 cells), lower library complexities (14,681) and a smaller fraction of reads in peaks per cell (30.0%) (**Fig. S4g,h**).

**Figure 2.**
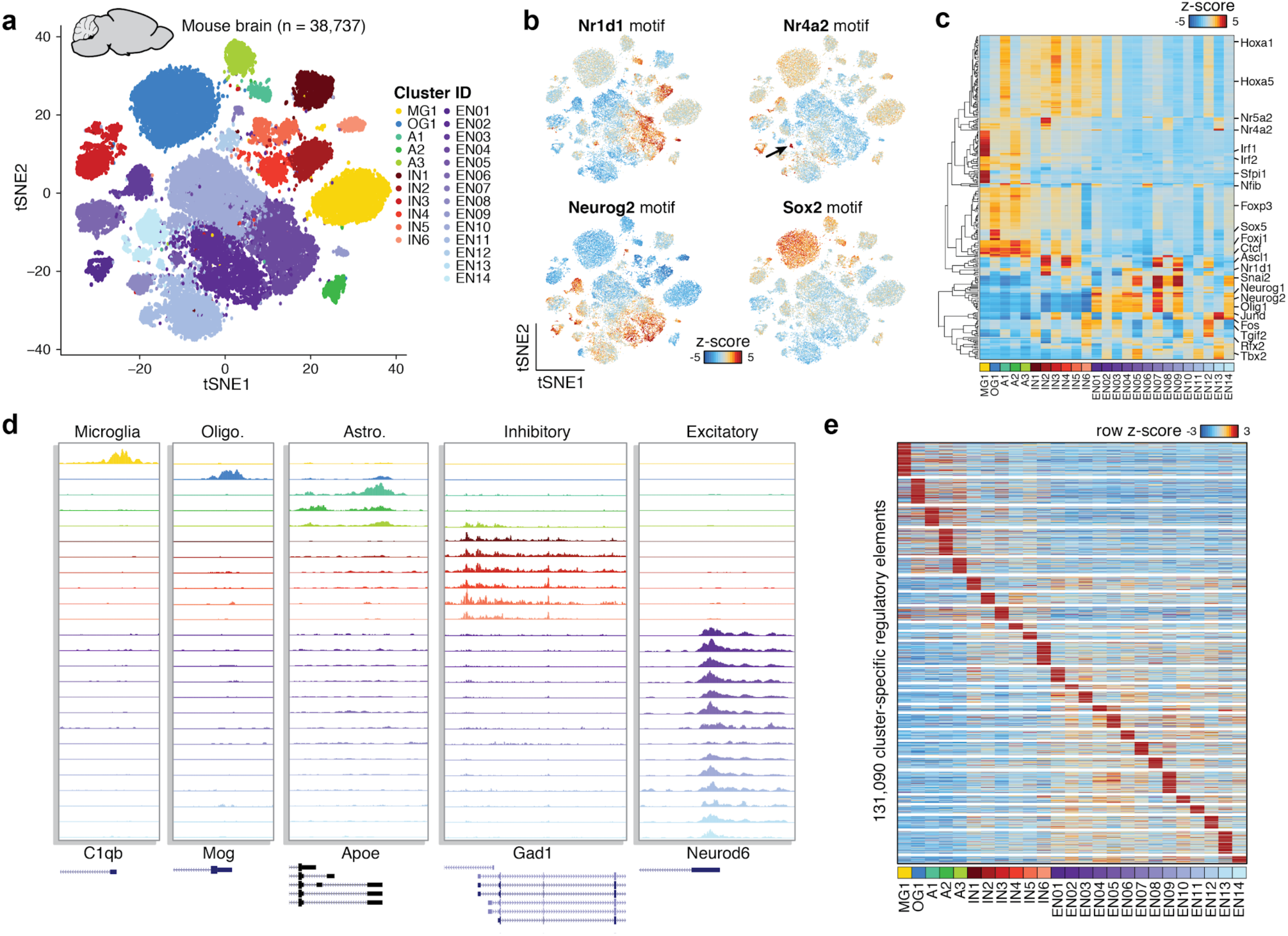
*De novo* classification of cell types in the mouse brain. (**a**) A t-SNE visualization of 38,737 cells derived from two mouse whole brains across 10 experimental replicates. Cells are colored by their identity across 25 clusters. (**b**) Cluster-specific activity of known TF regulators in the mouse brain, each panel depicts the chromVAR deviation score for each transcription factor motif. (**c**) Hierarchical clustering of the average per-cluster transcription factor deviation scores; relevant transcription factor motifs are highlighted. (**d**) Aggregate chromatin accessibility profiles per cluster surrounding the promoter region of known marker genes. (**e**) Chromatin accessibility signal across 165,968 cell type-specific peaks within clusters defined in the mouse brain.

To annotate these clusters, we used TF motif deviation scores (**Fig. 2b-c** and **Fig. S5a**) and promoter region accessibility scores (weighted-sum of chromatin accessibility ±100 kb around the TSS) at known cell type specific gene markers (**Fig. 2d** and **Fig. S5b-d**) to classify these clusters into the major cell types found in the brain. These clusters include microglia (MG1), oligodendrocytes (OG1), astrocytes (A1, A2, A3), inhibitory neurons (IN1-IN6) and excitatory neurons (EN01-EN14) (**Fig. 2a** and **Fig. S5**). Interestingly, we also identified an enrichment of the transcription factor motif Nr4a2 in EN13 (**Fig. 2b**) suggesting that this cluster corresponds to dopaminergic neurons, given the critical role of this TF in the development and maintenance of the dopaminergic system^28^. While this approach provided broad classifications for annotation of cell types, further work is needed to define chromatin accessibility markers unique to each neural cell type. In addition to the inference of cell types and cell type-specific TF master regulators, our approach also enabled the unbiased identification of 131,090 cell type-specific chromatin accessibility peaks (**Fig. 2e**), which further validates the unique identity of each cell cluster and provides a general resource for defining regulatory elements to drive cell type-specific reporters in effort to better understand the mouse brain^29^. In summary, we find that the dscATAC-seq platform provides a powerful means for defining and annotating cell states while further identifying cell type-specific chromatin accessibility.

### Development of droplet-based sciATAC-seq for massive-scale single-cell studies

Although the dscATAC-seq approach can be scaled to generate data for large cell numbers by simply performing the experiment across many replicates, as shown above (**Fig. 2**), we reasoned that experiments at this scale would be cost-prohibitive, laborious, and potentially susceptible to technical confounders across batches. We therefore sought to combine this approach with combinatorial indexing^19,23^ to improve throughput and enable multiplexing of multiple samples in a given experiment. To achieve this, we developed a method for droplet-based sciATAC-seq (dsciATAC-seq), wherein we load Tn5 transposase with barcoded DNA adapters to add well-specific DNA barcodes to open chromatin. Following barcoded transposition, transposed cells are pooled and loaded at high density to co-encapsulate multiple barcoded cells with multiple beads in each droplet (**Fig. 3a,b**). Following droplet co-encapsulation of transposed cells with barcoded beads and PCR reagents, we perform library preparation using our standard approach.

**Figure 3.**
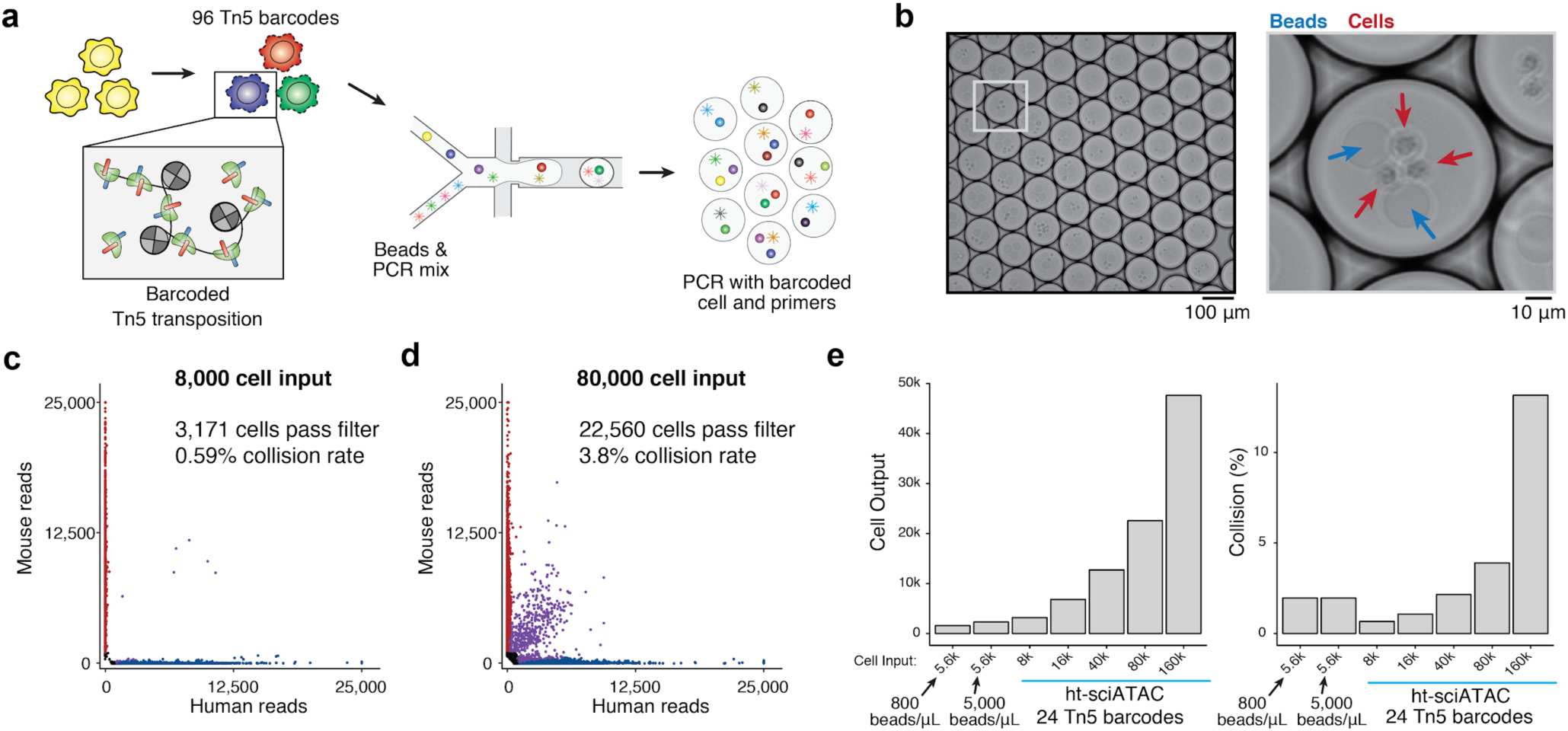
dsciATAC-seq enables massive scale single-cell experiments. (**a**) Schematic of dsciATAC-seq. Cells are transposed with barcoded Tn5, pooled, and then further processed through the droplet PCR microfluidic device. (**b**) Representative image of droplets containing multiple beads and cells. Blue arrows indicate beads and red arrows indicate transposed nuclei. (**c,d**) The number of unique reads aligning to the mouse or human genome from dsciATAC-seq profiles of human (K562) and mouse (3T3) cells with (**c**) 8,000 and (**d**) 80,000 cell input. (**e**) Summary of species mixing and cell yield results at variable cell inputs.

First implementing this technology with 24 transposase barcodes, we generated high-quality chromatin accessibility profiles for up to 50,000 cells in a single well of the device representing one experimental sample. Species mixing analysis confirmed that we could increase cell throughput approximately 10-fold while maintaining a collision rate lower than 5% using 24 transposase barcodes (**Fig. 3c-e** and **Fig. S6a**), and confirmed a further reduction in the overall detected collision rates at large cell inputs with 48 barcodes (**Fig. S6b**). Altogether, in this cell titration experiment, we generated 274,144 single-cell profiles demonstrating the massive scalability of this approach. Notably, to perform this experiment we separately purified and *in vitro* assembled barcoded Tn5^30^. As this proof-of-concept experiment did not utilize the optimized Tn5 described above, we observed fewer reads per cell, however maintained a high fraction of reads in peaks (72.2%). Together, these experiments demonstrate that barcoded Tn5 can enable super-Poisson loading of cells into droplets to achieve a significantly greater throughput for generating epigenomic profiles from 10^4^ to 10^5^ single-cells per experiment.

### Chromatin accessibility profiling of the native human bone marrow

Barcoded Tn5 transposition enables a significantly increased cell throughput and the opportunity to multiplex scATAC-seq to multiple conditions or samples. Notably, tissue-scale perturbations^31^ have been used to uncover diverse cell response dynamics^32^. We therefore reasoned that pooled stimulation across heterogenous cell types within bone marrow mononuclear cells (BMMCs) would provide unique avenues to understand the functional roles of epigenomic diversity within the human bone marrow. To achieve this, we used dsciATAC-seq using 96 transposase barcodes to profile BMMCs from two human donors before (untreated controls) and after stimulation, producing chromatin accessibility profiles for a total of 136,463 cells passing quality filters (**Fig. 4a** and **Fig. S7**).

**Figure 4.**
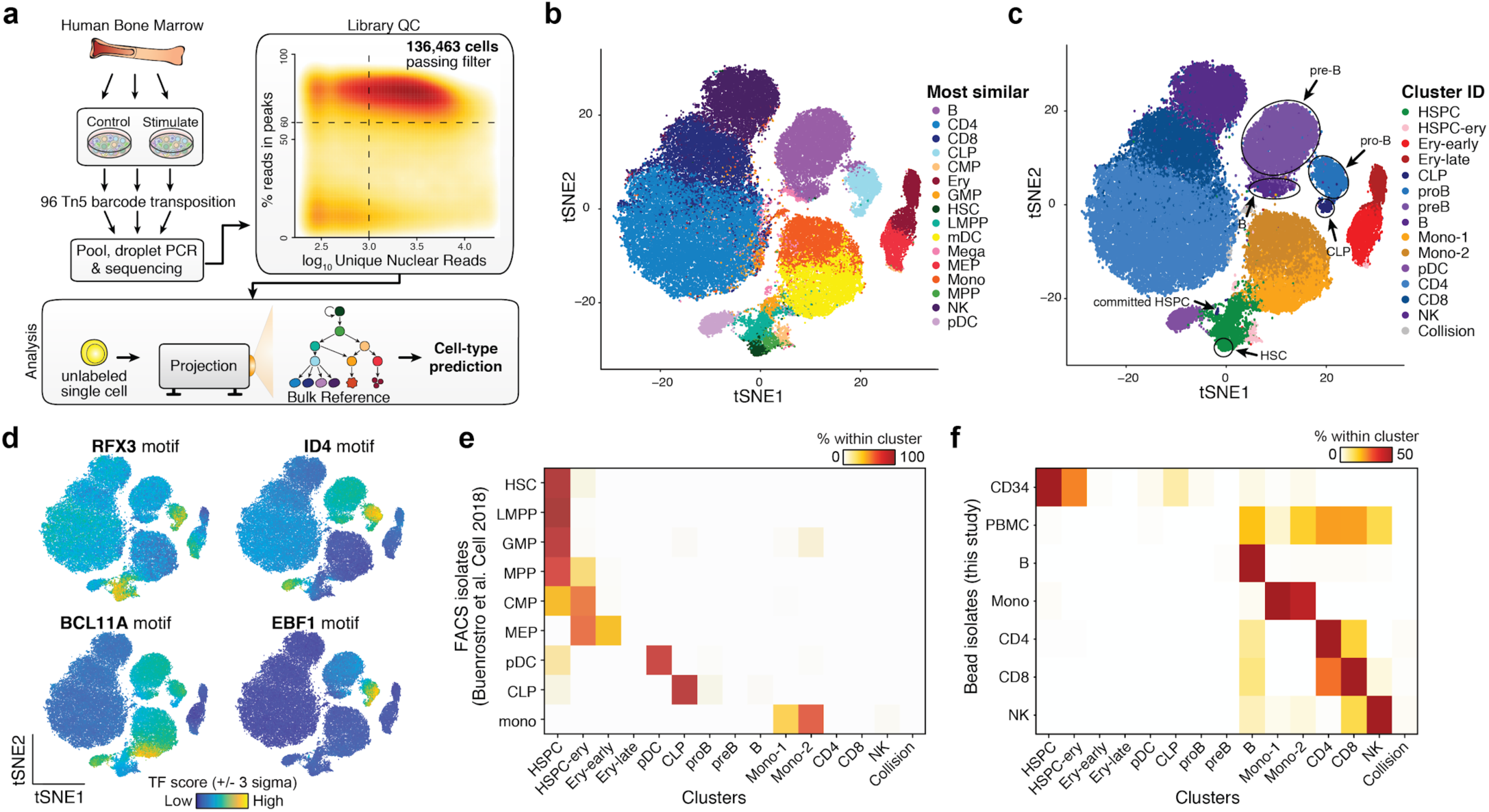
dsciATAC-seq of human bone marrow cells reveals the major lineages of hematopoietic differentiation. **(a)** Schematic of experimental and computational workflow used to assess bone marrow mononuclear cells dsciATAC-seq data. 96 Tn5 transposase barcodes are used to define different donors and stimulation conditions. Library QC box displays the summary of data passing quality filters across all assayed cells. 136,463 cells were identified passing filters of 60% reads in peaks and 1,000 unique nuclear reads. (**b,c**) Two dimensional t-SNE embedding of single bone marrow mononuclear cells without stimulation (*n*=60,495 cells). Cells are colored by (**b**) the most correlated cell type from a bulk ATAC-seq reference or (**c**) 15 *de novo* defined cluster assignments covering known hematopoietic cell types. Cell types covering the B-cell differentiation trajectory are highlighted. (**d**) Single-cells are colored by TF motif accessibility scores, computed using chromVAR^26^, for the motifs RFX3, ID4, BCL11A and EBF1. (**e,f**) Confusion matrix showing the percent overlap between (**e**) published scATAC-seq data^21^ and (**f**) isolated subsets collected in this study.

The reference map of 60,495 resting cells (untreated controls) revealed the major lineages of hematopoietic differentiation *de novo*. To analyze these reference datasets, we projected the untreated BMMCs onto hematopoietic development trajectories using a reference-guided approach, whereby single-cells are scored by principal components trained on bulk sorted hematopoietic ATAC-seq profiles^21^ (**Fig. 4a**) (see Methods). With this approach, we are able to visualize and predict cell labels given the bulk reference map of epigenomic states (**Fig. 4b**). Further, using the Louvain modularity method, we identified 15 distinct clusters from the 60,495 resting cells, which recapitulate the major constitutive cell types in the human hematopoietic system (**Fig. 4c**). These *de novo* derived single cell clusters reflect changes in chromatin accessibility mediated by key lineage-specific transcription factor motifs, including those associated with the step-wise progression of B cell development from hematopoietic stem cells (HSCs) to mature B cells (**Fig. 4c,d**). Furthermore, we observed unexpected epigenomic heterogeneity across transcription factor motifs, including CEBPD and BCL11A, within monocyte clusters (Mono-1 and Mono-2), which likely reflects the heterogenous developmental transitions from myeloid progenitors to mature monocytes, myeloid dendritic cells (mDCs) and granulocytes (**Fig. S8a**).

To validate the clusters and cell type annotations from our approach, we assigned previously profiled FACS-sorted single-cell ATAC-seq profiles from progenitors in human bone marrow and peripheral blood monocytes (2,034 cells)^21^ to clusters defined here. We classified these published single-cell data to clusters based on the minimum Euclidean distance of a single-cell profile to a cluster medoid. We observed significant overlap between each isolated subset and its corresponding cluster in the dsciATAC-seq data, validating the progenitor cell type annotations (**Fig. 4e**). Furthermore, we performed dscATAC-seq on CD34+ bone marrow progenitor cells, peripheral blood mononuclear cells (PBMCs), and from bead-enriched subpopulations from PBMCs to derive a total of 52,873 cells, which validated our cluster label assignments for mature cell types (**Fig. 4f** and **Fig. S8b**). We also used an orthogonal approach to visually validate these findings by dimensionality reduction using the uniform manifold projection (UMAP) algorithm^33^, which allows for data to be projected onto the dsciATAC-seq base dimensionality (**Fig. S8c-f**). Collectively, we have used this approach to define a reference epigenomic atlas of cell states within hematopoietic cells in the human bone marrow, highlighting the applicability of our combinatorial approach to generate accurate large-scale epigenomic maps to define cell types within primary human tissues.

### Regulatory consequences of multi-lineage stimulation

Our multiplexed, droplet-based sciATAC-seq method further provides a unique opportunity to decipher regulatory consequences of perturbation without concerns for batch-effects confounding experimental results. To characterize the response of each immune cell cluster to stimulation conditions, we explored the differences between our untreated control cells and *ex vivo* cultured and LPS-stimulated BMMCs (**Fig. 4a**). To determine *trans-*acting regulators altered in response to these perturbations, we developed an analytical strategy wherein we compute differential TF scores by i) defining a K-nearest neighbor map connecting stimulus to control conditions, and ii) computing differential TF scores by calculating the difference in TF scores between each cell and the average of 20-nearest stimulus cells (**Fig. 5a**). Interestingly, we found significant and highly correlated epigenomic responses to both *ex vivo* culture and LPS stimulation (**Fig. S9a-e**), suggesting that the effects of *ex vivo* culture dominates those induced by LPS. For clarity we simply refer to these conditions as “stimulation” for downstream analysis. With this stimulation data representing the full spectrum of bone marrow hematopoietic cell states, we found cell type-specific induction of a diverse repertoire of TF motifs (**Fig. 5b-d** and **Fig. S9f-j**). This list of differential TFs included induction of the Jun and NFkB motifs, largely localized to human hematopoietic stem and progenitor cells (HSPCs) (**Fig. 5b,c**), depletion of the SPIB motif in myeloid cell types (**Fig. 5d**) and relatively weak induction of MAFF (myeloid) and IRF8 motifs (MEP and CLP to pre-B) (**Fig. S9i,j**). Interestingly, Jun and NFkB were largely correlated in HSPCs, with the exception of CLP and early erythroid differentiation, wherein cells appeared to respond exclusively by NFkB motif induction (**Fig. 5b**).

**Figure 5.**
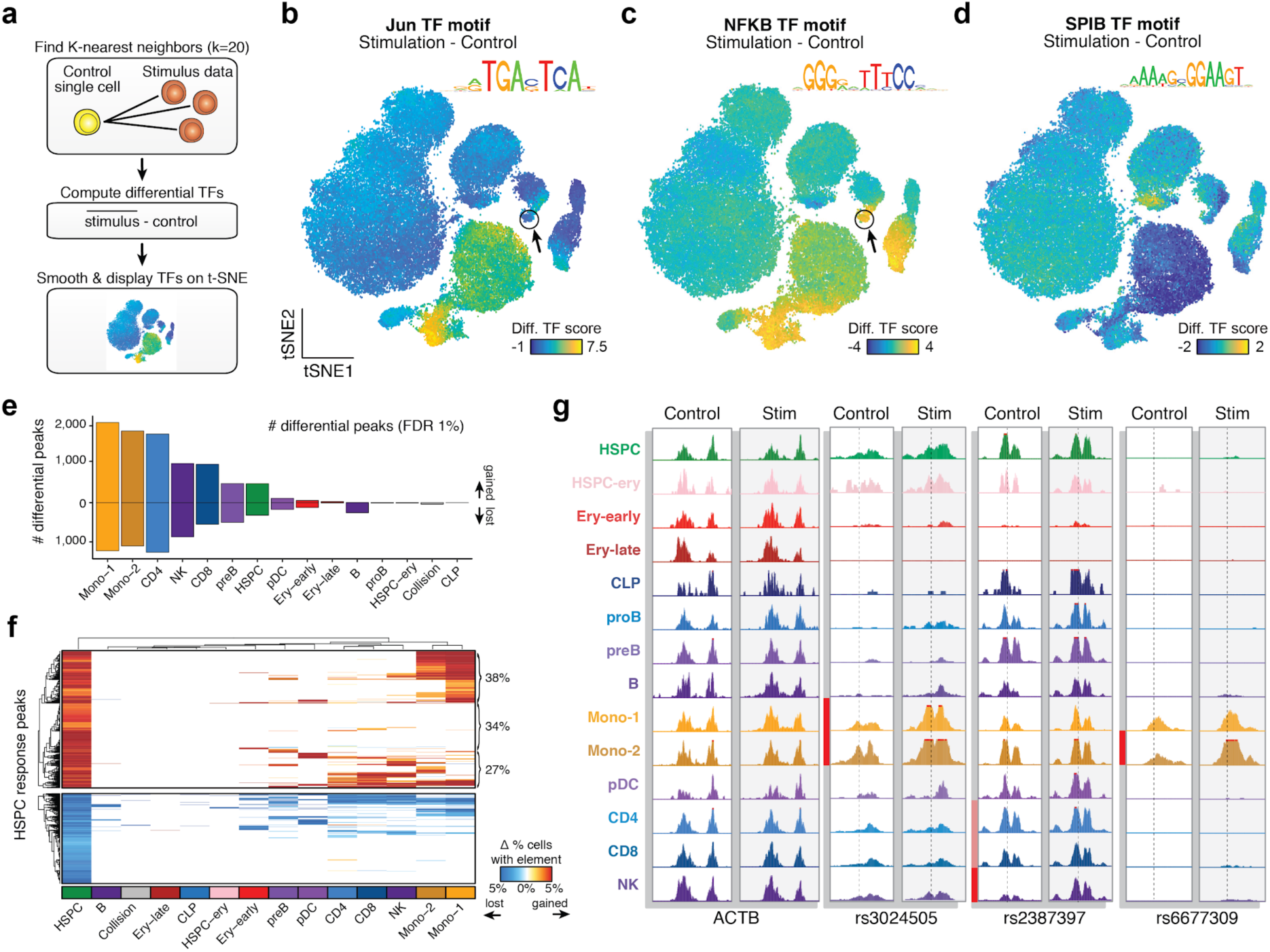
Identification of stimulus-response regulators in human bone marrow. (**a**) Schematic depicting the computational workflow for comparing stimulus versus control single-cell data. (**b-d**) Differential TF deviation scores for Jun, (**c**) NF-κB and (**d**) SPIB motifs in response to stimulation. (**e**) Summary of the number of differential chromatin accessibility peaks across each cluster at a false discovery rate (FDR) of 1%. Bars above the zero line represent gained chromatin accessibility peaks, bars below the zero line represent lost chromatin accessibility peaks. (**f**) Hierarchical clustering of peaks (top) gained or (bottom) lost across clusters, restricted to the differential peaks identified in HSPCs. (**g**) Locus specific views of the *ACTB* promoter and three fine-mapped variants identified through genome-wide association studies. The dotted line represents the location of the SNP in each window, the red bars (*p*<10^-5^) and pink bar (*p*<10^-1^) represent the statistical significance of the change in chromatin accessibility for each cell type cluster.

Next, we examined the *cis-*regulatory consequences of stimulus across our multi-lineage defined cell states. To compute differential chromatin accessibility peaks within each cluster, we devised a permutation test per peak, permuting control and perturbation cell labels, which allowed us to improve the robustness of our statistical methods by considering each cell as an independent observation (**Fig. S9k-l**; see Methods). This analysis revealed a total of 9,638 stimulus-responsive chromatin accessibility peaks (FDR 1%, **Table S4**). Interestingly, we broadly observed a gain in the total number of chromatin accessibility peaks, represented by the Mono-1 cluster with 2,114 peaks gained compared to 1,264 peaks lost (binomial p-value < 2.2e-16) (**Fig. 5e**). A global gain in chromatin accessibility upon stimulation was also corroborated by an approximate 20% gain in the average library complexity per-cell. The most prominent cell types that responded to the stimulation treatment included the two monocyte and CD4 T-cell clusters. Unexpectedly, we also observed 501 chromatin accessibility peaks gained in the HSPC cluster, and approximately 34% of these HSPC gained peaks were unique to HSPCs (**Fig. 5f**), thus uncovering an HSPC-specific stimulus response signature. Altogether, considering the TF motif and peak-specific analyses, we find that HSPCs respond to stimulus using the NFkB and Jun TF motifs to drive an HSPC-specific stimulus response. This finding supports reports suggesting that HSPCs are responsive to interferon-mediated immune signaling^34,35^, and may be used to further characterize the regulatory basis of interferon signaling in HSPCs to nominate chemical inhibitors to facilitate *ex vivo* expansion and gene editing of HSCs^36^ for hematopoietic stem cell transplantation (HSCT).

We further hypothesized that this approach to uncover cell type-specific stimulation changes could elucidate mechanisms of relevant cell types and regulatory regions for variants implicated in genome-wide association studies^37,38^. Towards this effort, we observed stimulation response chromatin accessibility peaks near the IL-10 locus in monocytes overlapping the pleiotropic rs302405 variant locus associated with Type 1 Diabetes (posterior probability (PP) = 0.38), Crohn’s Disease (PP=0.40), and Ulcerative Colitis (PP=0.41), as well as chromatin accessibility gains at the variant rs2387397 associated with Celiac Disease (PP=0.32) within the natural killer (NK) and T-cell clusters (**Fig. 5g**). Additionally, we observed a Mono-2 stimulation-specific peak overlapping rs6677309, a fine-mapped variant associated with multiple sclerosis (PP=0.49), near the *CD58* locus (**Fig. 5g**). Interestingly, CD58 presentation by activated monocytes has been shown to expand CD56+ NK cells^39^, which may promote an autoimmune response in multiple sclerosis^40^. Overall, this single experiment comprising 60,495 resting and 75,968 stimulated cells enabled the unbiased discovery of regulatory changes across stages of hematopoietic differentiation, and the unbiased identification of the regulatory consequences of *ex vivo* perturbation across multiple lineages, providing new opportunities to better define cell types within complex tissues and their relationship to stem cell therapy and autoimmune disease.

## Discussion

In the genomics era of cell atlases, a major goal of single-cell methods is to provide an unbiased classification of cell types and the epigenomic, transcriptomic and proteomic features that define them^41^. We find that scATAC-seq maps can provide information rich measurements of cells (10^5^ fragments per cell), which enables the identification of cell types and their underlying regulatory elements. Further, previous work has suggested that regulatory element activity may be a more accurate reflection of cell potential and perhaps more cell type-specific than gene expression measurements^17^. The scATAC-seq approach described here produces single-cell profiles at higher-throughput, improved yield and higher sequencing efficiency than previous scATAC-seq methods, providing a robust platform for identifying new cell types within heterogeneous tissues. We expect that the combination of this scATAC-seq approach with scRNA-seq profiling approaches will provide a more accurate definition of cell types and further integration of these data^21,27,42^ will enable opportunities to define mechanistic gene regulatory models to understand their function.

We present a series of technological innovations leading to a high-throughput epigenomic profiling approach that enables super-Poisson loading of cells and beads into microfluidic droplets. To achieve this we have developed a computational approach to identify droplets with multiple barcoded beads and paired this approach with combinatorial indexing by barcoded transposition to add multiple cells per droplet. Combining these approaches dramatically improves cell throughput to approximately ∼25,000 cells per well (100,000 cells per droplet device), which we expect may be further improved with optimizations of the approach and additional Tn5 barcodes. More generally, we expect this conceptual framework of combinatorial indexing coupled with a microfluidics device may be compatible with other methods for high-throughput PCR (e.g. microwells^43^) and other single-cell genomics assays leveraging combinatorial indexing for cell barcoding^44–46^.

This approach allows for multiplexing of many samples in a single experiment. In this work, we multiplex control and perturbation conditions across an entire tissue, enabling us to define shared and cell type-specific regulatory changes induced upon stimulation across diverse cell types. These advances for multiplexing experiments along with advances in high-throughput sequencing, opens new opportunities to define not only cell type-specific chromatin accessibility, but also changes across diverse genetic and environmental conditions. As such we expect this approach to be used to profile epigenomic variation across healthy individuals or from cohorts of diseased patients to determine the functional roles of both regulatory elements and cell types underlying common traits or in disease^47^. Altogether, these advances enable a new era of single-cell epigenomic studies at a massive scale, providing a powerful new tool to connect the vast repertoire of DNA regulatory elements to function.

## Acknowledgements

We thank members of the Buenrostro lab for useful discussions and critical assessment of this work. We thank Dan Norton at Bio-Rad, the SureCell ATAC-Seq Library Prep Kit product manager, for enabling the collaboration with the Broad Institute. Other Bio-Rad colleagues that we would also like to acknowledge for establishing and providing droplet related consumables are Duc Do, Bin Zhang, Preeti Pattamatta, Sean Cater, Lucas Frenz, Doug Greiner, and Jeremy Agresti. We also recognize Lena Christiansen, Allison Yunghans, and Lisa Watson from the Illumina Assay Development team, in addition to Fan Zhang and Felix Schlesinger at Illumina for their bioinformatics contributions. We are grateful to the Zhang lab for providing the Tn5 for combinatorial experiments. JDB, CAL, FMD and VKK acknowledge support by the Allen Distinguished Investigator Program, through The Paul G. Allen Frontiers Group. This work was further supported by the Chan Zuckerberg Initiative (CZI). CAL is supported by an NIH F31 grant (F31CA232670).

## Author contributions

FMD, JGC and ASK generated the data. CAL, VKK, and ZDB analyzed the data. FJS proposed the droplet scATAC-seq approach and oversaw the proof-of-concept studies performed by DP. MJA assisted in the development of computational resources. CAL, FMD and JDB wrote the manuscript with input from all authors. RL and JDB jointly supervised this work.

## Supplementary materials

Figure S1: Optimization of Tn5 transposition for dscATAC-seq.

Figure S2: Validation of bead merging computational approach.

Figure S3: Additional quality controls of dscATAC-seq.

Figure S4: Quality control information for the dscATAC-seq mouse brain dataset and comparison with existing data.

Figure S5: Chromatin accessibility scores for validation of cell clusters from mouse brain.

Figure S6: Species mixing analysis of dsciATAC-seq.

Figure S7: Quality control analysis of human bone marrow dsciATAC-seq data.

Figure S8: Cell types identified in the human bone marrow dsciATAC-seq data.

Figure S9: Stimulation of human bone marrow derived cells.

TableS1: Oligonucleotides used in this study.

TableS2: Human peripheral blood and bone marrow cells donor information.

TableS3: Cluster specific peaks identified in the mouse brain.

TableS4: Stimulus responsive chromatin accessibility peaks.

## Methods

### Cell lines

GM12878 (Coriell Institute for Medical Research) human lymphoblastoid cells were maintained in RPMI 1640 medium modified to include 2 mM L-glutamine (ATCC), 15% FBS (ATCC) and 1% Penicillin Streptomycin (Pen/Strep) (ATCC). K562 (ATCC) human chronic myelogenous leukemia cells were maintained in Iscove’s Modified Dulbecco’s Medium (IMDM) (ATCC) supplemented with 10% FBS and 1% Pen/Strep. NIH/3T3 (ATCC) mouse embryonic fibroblast cells were maintained in Dulbecco’s Modified Eagle’s Medium (DMEM) (ATCC) supplemented with 10% Calf Bovine Serum and 1% Pen/Strep. All cell lines were maintained at 37°C and 5% CO_2_ at recommended density and were harvested at mid-log phase for all experiments. All suspension cells were harvested using standard cell culture procedure, and adherent cells were detached using TrypLE Express Enzyme (Gibco). After harvesting, cells were washed twice with ice cold 1x PBS (Gibco) supplemented with 0.1% BSA (MilliporeSigma). Cells were then filtered with a 35 μm cell strainer (Corning) and cell viability and concentration were measured with trypan blue on the TC20 Automated Cell Counter (Bio-Rad). Cell viability was greater than 90% for all samples.

### Mouse tissues

Flash frozen adult mouse whole brain tissue was purchased from BrainBits (SKU: C57AWB). Nuclei isolation was performed using the Omni-ATAC protocol for isolation of nuclei from frozen tissues^22^. Nuclei permeability and concentration were measured with trypan blue on the TC20 Automated Cell Counter. For all samples, over 95% of the nuclei were permeable to trypan blue, meaning that the nuclei isolation was successful.

### Human peripheral blood and bone marrow cells

Cryopreserved human bone marrow (BM) mononuclear cells, isolated BM CD34+ stem/progenitor cells, peripheral blood mononuclear cells (PBMC), and isolated peripheral blood CD4+, CD8+, CD14+, CD19+ and CD56+ cells were purchased from Allcells (see **Table S2** for catalog numbers and donor information). Cells were quickly thawed in a 37°C water bath, rinsed with culture medium (IMDM medium supplemented with 10% FBS and 1% Pen/Strep) and then treated with 0.2 U/μL DNase I (Thermo Fisher Scientific) in 10 mL of culture medium at 37°C for 30 min. After DNase I treatment, cells were washed with medium once and then twice with ice cold 1x PBS + 0.1% BSA. Cells were then filtered with a 35 μm cell strainer (Corning) and cell viability and concentration were measured with trypan blue on the TC20 Automated Cell Counter (Bio-Rad). Cell viability was greater than 80% for all samples.

### Human bone marrow mononuclear cells stimulations

BM mononuclear cells were quickly thawed in a 37°C water bath, rinsed with culture medium (RPMI 1640 medium supplemented with 15% FBS and 1% Pen/Strep) and then treated with 0.2 U/μL DNase I in 10 mL of culture medium at 37°C for 30 min. After DNase I treatment, cells were washed with medium once, filtered with a 35 μm cell strainer and cell viability and concentration were measured with trypan blue on the TC20 Automated Cell Counter. Cell viability was greater than 90% for all samples. Cells were plated at a concentration of 1 × 10^6^ cell/mL, rested at 37°C and 5% CO_2_ for 1 h and then either incubated in serum containing media (RPMI 1640 medium supplemented with 15% FBS and 1% Pen/Strep) at 37°C and 5% CO_2_ for 6 h (ex vivo culture) or treated with 20 ng/mL Lipopolysaccharide (LPS) (tlrl-3pelps, Invivogen) for 6 h (LPS stimulation). After stimulation, cells were washed twice with ice cold 1x PBS + 0.1% BSA and cell viability and concentration were measured with trypan blue on the TC20 Automated Cell Counter. As a control, we processed cells immediately after counting, without any incubation.

### Cell lysis and tagmentation

For a detailed description of tagmentation protocols and buffer formulations refer to the SureCell ATAC-Seq Library Prep Kit User Guide (17004620, Bio-Rad). Harvested cells and tagmentation related buffers were chilled on ice. For cell lines, a protocol based on Omni-ATAC was followed^22^. Briefly, washed and pelleted cells were lysed with the Omni-ATAC lysis buffer containing 0.1% NP-40, 0.1% Tween-20, 0.01% Digitonin, 10 mM NaCl, 3 mM MgCl_2_, and 10 mM Tris-HCl pH7.4 for 3 min on ice. The lysis buffer was diluted with ATAC-Tween buffer that only contains 0.1% Tween-20 as a detergent. Cells were collected and resuspended in OMNI Tagmentation Mix containing ATAC Tagmentation Buffer and ATAC Tagmentation Enzyme that are parts of the SureCell ATAC-Seq Library Prep Kit (17004620, Bio-Rad). The OMNI Tagmentation Mix was buffered with 1X PBS supplemented with 0.1% BSA. Cells were mixed and agitated on a ThermoMixer (5382000023, Eppendorf) for 30 min at 37°C. Tagmented cells were kept on ice prior to encapsulation.

For PBMCs and BM mononuclear cells, lysis was performed simultaneously with tagmentation. Washed and pelleted cells were resuspended in Whole Cell Tagmentation Mix containing 0.1% Tween-20, 0.01% Digitonin, 1X PBS supplemented with 0.1% BSA, ATAC Tagmentation Buffer and ATAC Tagmentation Enzyme. Cells were tagmented using a thermal protocol and maintained thereafter as described in the Omni-ATAC protocol described above.

For mouse tissues, nuclei were washed with ATAC-Tween buffer containing 0.1% Tween-20, 10 mM NaCl, 3 mM MgCl_2_, and 10 mM Tris-HCl pH7.4 prior to the whole cell protocol described above.

### Droplet library preparation and sequencing

For a detailed protocol and complete formulations, refer to the SureCell ATAC-Seq Library Prep Kit User Guide (17004620, Bio-Rad). Tagmented cells or nuclei were loaded onto a ddSEQ Single-Cell Isolator (12004336, Bio-Rad). Single-cell ATAC-seq libraries were prepared using the SureCell ATAC-Seq Library Prep Kit (17004620, Bio-Rad) and SureCell ddSEQ Index Kit (12009360, Bio-Rad). Bead barcoding and sample indexing were performed in a C1000 Touch™ Thermal cycler with a 96-Deep Well Reaction Module (1851197, Bio-Rad): 37°C for 30 min, 85°C for 10 min, 72°C for 5 min, 98°C for 30 sec, 8 cycles of 98°C for 10 sec, 55°C for 30 sec, and 72°C for 60 sec, and a single 72°C extension for 5 min to finish. Emulsions were broken and products cleaned up using Ampure XP beads (A63880, Beckman Coulter). Barcoded amplicons were further amplified using a C1000 Touch™ Thermal cycler with a 96-Deep Well Reaction Module: 98°C for 30 sec, 6-9 cycles (cycle number depending on the cell input, Section 4 Table 3 of the User Guide) of 98°C for 10 sec, 55°C for 30 sec, and 72°C for 60 sec, and a single 72°C extension for 5 min to finish. PCR products were purified using Ampure XP beads and quantified on an Agilent Bioanalyzer (G2939BA, Agilent) using the High-Sensitivity DNA kit (5067-4626, Agilent). Libraries were loaded at 1.5 pM on a NextSeq 550 (SY-415-1002, Illumina) using the NextSeq High Output Kit (150 cycles; 20024907, Illumina) and sequencing was performed using the following read protocol: Read 1 118 cycles, i7 index read 8 cycles, and Read 2 40 cycles. A custom sequencing primer is required for Read 1 (16005986, Bio-Rad; included in the kit).

### dsciATAC-seq methods

#### Assembly of indexed Tn5 transposome complexes

To generate indexed Tn5 transposome complexes, we modified the Illumina Nextera Read 1 Adapter to contain a 6 nt barcode (96 distinct barcodes, see **Table S1** for barcode sequences). Each indexed oligo was mixed with the Illumina Nextera Read 2 Adapter and annealed to a 15 nt mosaic end complementary oligonucleotide (5’ phosphorylated and 3’ Dideoxy-C) (**Table S1**). All oligonucleotides were HPLC purified (IDT). For the annealing reaction, oligonucleotides were mixed at a 1:1:2 molar ratio (Read 1: Read 2: complementary mosaic end) at 100 μM final concentration in 50mM NaCl. The mixture was incubated at 85°C, ramped down to 20°C at a rate of −1°C/min, and then 20°C for 2 additional minutes. After being diluted 1:1 in glycerol, the annealed oligonucleotide mixture was then mixed 1:1 with 14.8 μM purified Tn5 (Tn5 purified as previously described^30^). The Tn5/oligonucleotide mixture was incubated for 30 min at room temperature and then kept at −20°C prior to the tagmentation reactions.

#### Species mixing controls

Human and mouse cell lines were processed and lysed using the Omni-ATAC-seq protocol as described above. For the 24-plex control experiment in Figure 3 and S6, K562 and NIH/3T3 cells were mixed at a 1:1 ratio and tagmented with Tn5 loaded with indexed oligonucleotides 1-3, 13-15, 25-27, 37-39, 49-51, 61-63, 73-75, 85-87 (**Table S1**) in 50 μL reactions (10 μL of indexed Tn5 per reaction) with 25,000 cells each. Cell line tagmentation buffer components and reaction conditions were the same as described above. After the tagmentation reaction, all cells were pooled, washed with tagmentation buffer without Tn5 and processed using our standard protocol for droplet library preparation and sequencing. Different cell numbers were used as input, as indicated in **Fig. 3** and **S6**.

For the 48-plex control experiment in Figure S6, K562 and NIH/3T3 cells were mixed at a 1:1 ratio and tagmented with Tn5 loaded with indexed oligonucleotides 1-6, 13-18, 25-30, 37-42, 49-54, 61-66, 73-78, 85-90 (**Table S1**) in 50 μL reactions (10 μL of indexed Tn5 per reaction) with 25,000 cells each. Cell line tagmentation buffer components and reaction conditions were the same as described above. After the tagmentation reaction, all cells were pooled, washed with tagmentation buffer without Tn5 and processed using our standard protocol for droplet library preparation and sequencing. Different cell numbers were used as input, as indicated in **Fig. S6**.

#### Human BM mononuclear cells stimulations

BM-MNCs from 2 donors were stimulated and washed as described above. For the experiment in Figures 4 and 5, BM-MNCs were tagmented with Tn5 loaded with indexed oligonucleotides 1-96 in 20 μL reactions (4 μL of indexed Tn5 per reaction) with 8,000 cells each (Control, ex vivo culture and LPS stimulation as described above). BM-MNC tagmentation buffer components and reaction conditions were the same as described above. After the tagmentation reaction, all cells were pooled, washed with tagmentation buffer without Tn5 and processed using our standard protocol for droplet library preparation and sequencing. Pooled cells were split into 16 different samples for droplet library preparation, with varying cells inputs (20,000, 40,000 or 80,000 cells). After sequencing, data from all 16 samples were merged for the analyses.

Sequencing data for the dsciATAC-seq experiments were processed with bap as described below using the “--tn5-aware” flag that inhibits cell merging across different Tn5 barcodes.

### Bioinformatic pre-processing

#### Raw read processing

Per-read bead barcodes were parsed and trimmed using UMI-TOOLs (https://github.com/CGATOxford/UMI-tools)^48^, and the remaining read fragments were aligned using BWA (http://bio-bwa.sourceforge.net/) on the Illumina BaseSpace online application. Constitutive elements of the bead barcodes were assigned to the closest known sequence allowing for up to 1 mismatch per 6-mer or 7-mer (mean >99% parsing efficiency across experiments). For the dsciATAC-seq experiments, bead barcodes were parsed using a custom python script aware of the 96 possible Tn5 barcodes. All experiments were aligned to the hg19 or mm10 reference genomes (or a combined reference genome in the case of species mixing experiments).

To identify systematic biases (i.e. reads aligning to an inordinately large number of barcodes), barcode-aware deduplicate reads, and perform bead merging (see below), we developed a software suite called the **b**ead-based **A**TAC-seq **p**rocessing (bap) tool. This software uses as input a .bam file for a given experiment with a bead barcode identifier indicated by a SAM tag. We generalized this pre-processing pipeline to handle other datasets (Fluidigm C1, sciATAC-seq) to enable consistent comparisons across various technologies (**Fig. 1**).

#### Knee-calling

To identify high-quality cells, knee calling was performed by first generating a Gaussian kernel density estimate (KDE) of the log10 transformed unique nuclear read counts per bead or droplet barcode. The KDE was then used to create a density distribution of 10,000 evenly spaced values between the minimum and maximum log10 transformed unique nuclear read counts. Local minima in this density distribution were identified and used as potential knee/inflection points below which barcodes were filtered from further analysis. This same procedure was applied to the bap Jaccard index for pairs of bead barcodes to identify beads originating from the same droplet.

#### Identification of multiple beads per droplet

An integral part of the technique described herein relies on the robust identification of pairs of bead barcodes that share exact insertions at a rate that exceeds what may be expected by chance. We note that our procedure readily enables multiple beads per droplet (**Fig. S2**). First, highly abundant barcodes are detected in the experiment wherein each unique barcode sequence is quantified among nuclear-mapping reads, and our knee calling algorithm establishes a per-experiment bead threshold. Next, sequencing reads assigned to a bead barcode passing filter are de-duplicated using the insert positions of the paired-end reads (as previously implemented in Picard tools).

After initial deduplication, we further remove paired-end reads that map to more than 6 bead barcodes, reasoning that these represent a technical confounder. Next, for each pair of bead barcodes passing the initial knee, we compute the Jaccard index over the insertion positions of reads, providing a measure of how similar the Tn5 insertions are between any pair of bead barcodes. From these pairwise Jaccard index statistics, we perform a second knee call to determine pairs likely to have originated from the same droplet (**Fig. S2d**). Finally, to assign droplet-level barcodes, we then loop over the original bead barcodes in order of their original nuclear read abundance. For a given bead barcode, if it is paired with any other bead barcodes that passed the pairwise knee, those bead barcodes are “merged” into one droplet barcode. This iteration repeats until all bead barcodes have been assigned to precisely one droplet barcode. To facilitate comparisons without droplet merging (e.g. **Fig. S2j,k**), our pipeline facilitates the “--one-to-one” flag, which maps one bead barcode onto one droplet barcode; this option was employed primarily to process other scATAC-seq datasets that would not have beads that would require merging.

#### Species mixing analysis

We carried out the same quantification procedure for all species mixing datasets analyzed in this work. Namely, reads were mapped to a hybrid hg19-mm10 reference genome using BWA. Cells were identified using the bap knee calling described above. The output of this pipeline yields the number of unique nuclear reads mapping to the mouse and human genomes, which were compared per-cell. We further excluded cells with less than 1,000 reads mapping to either the human or mouse genomes and identified collisions as those that had less than a 10x enrichment over the minor genome. The overall collision rate is reported as the number of annotated collision cells over the total number of cells compared (mouse + human + collisions).

#### Peak calling

For each scATAC-seq experimental sample, chromatin accessible summits were called using MACS2 callpeak with custom parameters previously described^17^. To generate a non-overlapping set of peaks per analysis, we first extended summits of each experiment to 500 bp windows (+/- 250 bp). We combined these 500 bp peaks, ranked them by their summit significance value, and retained specific non-overlapping peaks based on this ordering. We further removed peaks that overlapped the ENCODE blacklist and a custom mitochondrial blacklist generated by aligning a synthetic mtDNA genome to the nuclear genome (https://github.com/buenrostrolab/mitoblacklist).

#### Library complexity estimation

Per-cell library complexities were estimated using the Lander-Waterman equation^49^ using a custom R function translated from a previously established Java function implemented in Picard tools. Per-cell counts of total number of mapped nuclear reads passing quality filters and the number of unique nuclear reads served as inputs. The library complexity thus represents a metric that estimates the total number of unique nuclear reads given by the cell independent of sequencing depth.

### Comparison to public datasets

To benchmark the dscATAC-seq platform against existing datasets, we downloaded raw sequencing data (.fastq format) for GM12878 cells via three different combinatorial indexing scATAC-seq methods^19,24,25^ and 384 cells processed with the Fluidigm C1^18^ from GEO. All dataset were processed using the same pipeline, which included BWA alignment and downstream processing with bap using the “--one-to-one” flag that skips bead merging. We note that in all three combinatorial indexing scATAC-seq experiments, GM12878 cells were mixed with mouse cells. As such, we compared only annotated human cells (>9:1 ratio of human: mouse cells) from these experiments for downstream analysis.

To determine the correlation between single-cell ATAC-seq experiments, we used a merged peak set comprising of 175,581 combined DNase-seq hypersensitivity peaks from GM12878 and K562 made available through the ENCODE Project. The sum of single cells (agnostic to cell ID) were compared against bulk Dnase-seq profiles generated from ENCODE and Omni-ATAC^22^. To score the fraction of reads in peaks across single cell experiments, we used only the GM12878 DNase-seq peak set (124,321 peaks) to ensure that peak selection did not bias our quantification and comparison of technologies.

### Validation of multiple beads per droplet inference

To validate our ability to merge cells marked by multiple droplet beads, we introduced a diverse library of random oligonucleotides (14 nucleotides random region, see **Table S1** for full sequence) to our microfluidic reaction (**Fig. S2**). Human PBMCs were processed with this library of random oligos at bead concentrations of 200, 800, and 5,000 beads/μL, spanning the ranges used for the data presented in this work. The random oligonucleotides were spiked in to the cells at a final concentration of 5 nM after the tagmentation reaction, and samples were processed and sequenced using our standard protocol (described above). Among pairs of beads merged, the average number of oligos observed per bead ranged from 792-1,979.

We reasoned that bead barcodes sharing a noticeable overlap of these oligos (**Fig. S2a,b**) would be barcodes from two beads contained in the same droplet. We identified reads containing our random oligo by first identifying the 15 bp constant sequence and subsequently parsing the 14 bases downstream of the constant sequence (**Table S1**). For each experiment, we called a knee on the bead barcode pairwise Jaccard indices and computed the overlap of random sequences observed (**Fig. S2c**) for barcodes passing the nuclear read knee. For pairs of bead barcodes passing the oligo overlap knee, we annotated these as true positives.

Next, we computed our bap metric pairwise for each bead barcode using the overlap of pairs of inserts over each fragment (or paired-end read). This produces a metric for all pairs of bead barcodes with at least 500 unique nuclear reads observed per barcode (**Fig. S2d**). Using the true-positives defined from the random oligos data and a continuous overlap metric from bap, we computed precision-recall and receiver operating curves (mean area under the receiver-operating curve (AUROC) = 1.000 and mean area under the precision recall curve (AUPRC) = 0.997 (**Fig. S2g**)). We further compared other possible metrics for bead merging, including Pearson and Spearman correlation and a Jaccard index over reads in peaks, finding that our approach was the most robust and specific (**Fig. S2g**). We note that the library of random oligonucleotides provides a completely orthogonal measure of bead overlap compared to the nuclear DNA fragments used in the bap algorithm.

### Theory of beads and droplet concentrations

In this setting, we are interested in estimating the number of beads per droplet at variable bead concentrations using observed data. Given that our observed data does not yield any droplets with zero beads (cells not captured) nor can any measurement be relied on with greater than 6 beads (physical limit for beads in droplet; observed values likely reflect merged droplets), the observed number of beads per droplet is modeled by a double-truncated Poisson distribution. The probability density function of a double-truncated Poisson distribution for a single observation can be written as follows:

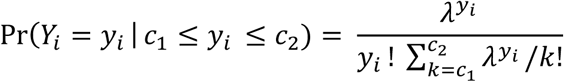

Here, *c*_1_ is our lower bound (1 in our case) of the empirical data and *c*_2_ is the upper bound (in our case 6) for observed numbers of beads / droplet *y*. Let *i* ∈ {1,2, …, *n*}. Then, we observe *n* cells and *y*_*i*_ denotes the number of beads per drop for cell *i*. The log likelihood (*l*) of observing a value can thus be computed as follows:

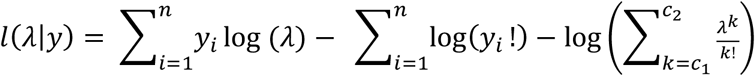

Here, a closed form solution of *λ* (parameter of the Poisson distribution indicating the mean number of beads per cell) is impossible. Thus, we estimate the value using the **optim()** function in R, providing the maximum likelihood estimate (MLE).

Given the MLE estimate for *λ*, from plugging into the Poisson PDF, we can trivially compute:

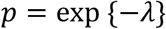

where *p* is the proportion of droplets with 0 beads. We can then approximate the number of droplets with a barcode as 1 – *p*. Empirical values of *λ* were determined using GM12878 data at different bead concentrations (800 and 5,000 beads/μL) and were found to be robust across the various datasets analyzed throughout.

### *De novo* K-mer clustering

Here, we present the use of a *de novo* k-mer feature-reduction approach agnostic to reduce the dimensionality of our scATAC-seq profiles. Specifically, we computed biased-corrected deviation z-scores for *K* k-mers and a set of *S* samples (scATAC-seq cells) with *P* peaks computed via the chromVAR methodology. Here, our implementation utilizes a binarized matrix ***M*** (dimension *P* by *S*) where *m*_*ik*_ is 1 if kmer *k* is present in peak *i* and 0 otherwise. For all applications, we used k = 7, resulting in (4^7^ / 2) 8,192 7-mers. Using the matrix of fragment counts in peaks ***X***, where *x*_*i,j*_ represents the number of fragments from peak *i* in sample *j*, a matrix multiplication ***X***^***T***^**. *M*** yields the total number of fragments for *S* samples (rows) and *K* 7-mers (columns). To compute a raw weighted accessibility deviation, we compute the expected number of fragments per peak per sample in ***E***, where *e*_*ij*_ is computed as the proportion of all fragments across all samples mapping to the specific peak multiplied by the total number of fragments in peaks for that sample:

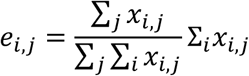

Analogously, ***X***^***T***^**. *E*** yields the expected number of fragments at peaks containing a given 7-mer sequence Using the ***M***, ***X***, and ***E*** matrices, we then compute the raw weighted accessibility deviation matrix ***Y*** for each sample *j* and k-mer *k* (*y*_*jk*_) as follows:

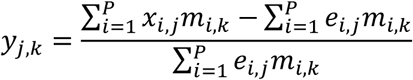

To correct for technical confounders present in assays (differential PCR amplification or variable Tn5 tagmentation conditions), we utilize the background peak sampling methodology previously described in chromVAR by generating a background set of peaks intrinsic to the set of epigenetic data examined. The matrix ***B***^(*b*)^ encodes this background peak mapping where 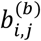 is 1 if peak *i* has peak *j* as its background peak in the *b* background set (*b* ∈ {1,2, …, 50}) and 0 otherwise. The matrices ***B***^(*b*)^**. *X*** and ***B***^(*b*)^**. *E*** thus give an intermediate for the observed and expected counts also of dimension *P* by *S*. For each background set *b*, sample *j*, and k-mer *k*, the elements 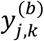 of the background weighted accessibility deviations matrix ***Y***^(*b*)^ are computed as follows:

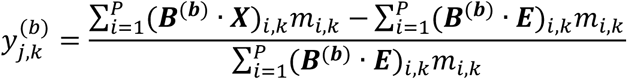

After the background deviations are computed over the 50 sets, the bias-corrected matrix ***Z*** for sample *j* and k-mer *k* (*z*_*jk*_) can be computed as follows:

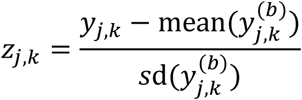

where the mean and variance of 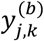 is taken over all values of *b* (*b* ∈ {1,2, …, 50}). As many of the 8,192 7-mers are highly correlated, we then use the top principal components of the matrix ***Z*** as input for downstream processes, including the Louvain clustering and t-SNE embedding.

### Cell type specific open chromatin peak identification in mouse brain

Here, we defined pseudo-bulk cell types by aggregating the total counts over each Louvain cluster. The peak x cell type count matrix was count-per-million (CPM) normalized, and peaks with an overall mean CPM > 1 were retained. This filtered peak x cell type matrix was then z-score transformed. We identified 131,090 cell-type specific chromatin accessibility peaks with a z-score > 3 in at least one cell type, which were assigned to clusters based on the maximum z-score value (**Fig. 2e**).

### Promoter region chromatin accessibility scores

To further validate our *de novo* clusters from the whole mouse brain, we computed per-cluster promoter region chromatin accessibility scores representing a weighted-sum of chromatin accessibility around the transcription start site (TSS) of each gene in our reference data. Specifically, for gene ***g*** and cluster *i*, we define a chromatin accessibility score ***g***_*i*_ from the following:

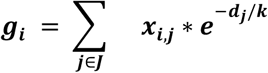

Here, ***x***_*i,j*_ represents the counts-per-million normalized chromatin accessibility count for cluster *i* and chromatin accessibility peak *j*. Accessibility peaks used per gene *J* were restricted to those within 100,000 bp of a corresponding TSS, and *d*_*j*_ represents the distance (in base pairs) between the TSS and the center of peak *j*. The scaling constant, *k*, was fixed to 5,000 for all chromatin accessibility score computations.

### Bulk-guided clustering

Bulk-guided clustering of single cells (**Fig. 4**) was performed as previously described^21^. Briefly, a matched peakset (*k*=156,311 peaks) was used for both BMMCs dsciATAC-seq (*n*=136,463 single cells), and bulk ATAC-seq profiles previously generated for sorted hematopoietic cell populations (16 cell types)^17,21,38^. PCA was first run on quantile-normalized bulk ATAC-seq data generating principal components (PCs) capturing variation across cell types. Single cells were then projected in the space of these bulk-trained PCs by multiplying the scATAC-seq reads in peaks matrix with the peaks x PC loading coefficients matrix to yield a matrix of single-cell projection scores (cells x PCs). The derived single-cell scores were then scaled and centered, and the corresponding single-cell data visualized using t-SNE. Predicted labels for single cells were obtained by correlating projected single cell scores with bulk PC scores, and choosing the most-correlated bulk cell type based on Pearson correlation coefficient. To define clusters for the control (unstimulated) BMMC dataset (**Fig. 4c**), Louvain clustering was performed using the igraph package where the 20 nearest neighbors per cell were used to build the embedding.

### Single-cell classification

To assign most-alike clusters generated form the 15 clusters of the control (unstimulated) BMMC dataset (**Fig. 4c**) to additional datasets (**Fig. 4e,f**), the medoids of each per-cluster principal component were determined over all cells assigned from the Louvain clustering at baseline. Next, for each cell from a new dataset (i.e. FACS-sorted populations and stimulation-response cells), we assigned to the cell a reference cluster based on the minimum Euclidean distance between each cell’s principal components and the medoids of the clusters.

### Analysis of differential TF motifs

To compute differential TF scores in normal and stimulation conditions, we determine the 20-nearest stimulus condition neighbors for each single-cell in the resting condition using the bulk-guided PC scores and a Pearson correlation distance metric. To calculate differential TF motifs, we subtract the mean of the 20 stimulus cells by the TF score for each cell in the normal condition. Last, to suppress noise in the comparison, we smooth the differential TFs by taking the mean of the 20 nearest neighbors in the control condition. Again, the nearest neighbors are calculated using the bulk-guided PC scores, with Pearson correlation as a distance metric.

### Differential peak identification in bone marrow stimulation

We devised a permutation test that accessed whether the proportion of cells with an accessibility element was differential between the stimulated and resting conditions, controlling for overall differences in accessibility (using measures at promoters). First, we filtered our consensus peak set such that the given peak was accessible in at least 1% of cells irrespective of stimulation or resting. Then, for an individual regulatory element *i*, we determined the proportion of cells in the resting *p*_*r*_ and the stimulated *p*_*s*_ conditions that observed one or more fragments overlapping the accessibility peak. Next, we computed the proportion of all promoters annotated in our dataset for both resting 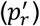 and stimulated 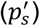. Our observed differential statistic thus is given by:

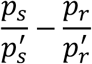

To determine statistical significance, we permuted the stimulation and resting labels 1,000 times to generate a permuted distribution. We observed the corresponding Z-statistic (**Fig. S9l**) to be centered with a largely-Gaussian distribution. After converting these Z-statistics to p-values, we computed a per-cluster false discovery rate (FDR) and established a significance threshold of 1% uniformly across clusters. We further computed an effect size of the difference between stimulated and resting, given simply by *p*_*r*_ − *p*_*s*_.

### Overlap with fine-mapped GWAS SNPs

To identify regulatory regions affected by our stimulation conditions that may be relevant for human disease, we overlapped differential peaks identified per cell type with single nucleotide polymorphisms (SNPs) identified through genetic fine-mapping studies of 21 immune traits as previously described^37^. Specifically, we downloaded the per-SNP meta-data available online (http://pubs.broadinstitute.org/pubs/finemapping/dataportal.php) and intersected differentially-accessible peaks with annotated positions of fine-mapped variants with a posterior probability > 0.3 computed by PICS^37^ across all reported traits.

## Supplemental Figures

**Figure S1.**
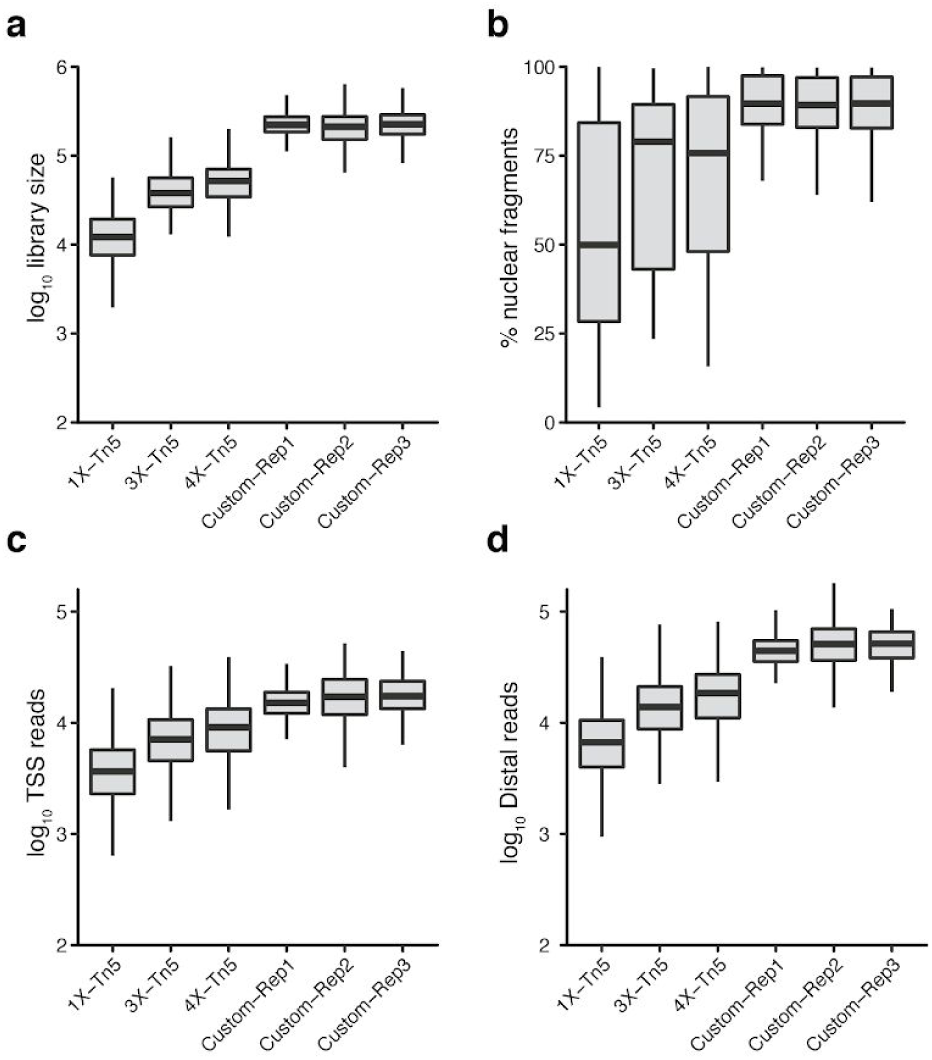
Optimization of Tn5 transposition for dscATAC-seq. (**a**) Per-cell library complexity estimated by the Lander-Waterman equation for variable Tn5 inputs. Different concentrations (1x, 3x, 4x) of commercial Tn5 are compared against 3 replicates of a custom Tn5 optimized for dscATAC-seq. (**b**) Fraction of reads mapping to the nuclear genome for each of the Tn5 concentrations. The remaining reads map to the mitochondrial genome. (**c**) Number of unique reads mapping near transcription start sites or (**d**) distal regulatory elements for the same Tn5 conditions. Center line, median; box limits, first and third quartiles; whiskers, 1.5x interquartile range.

**Figure S2.**
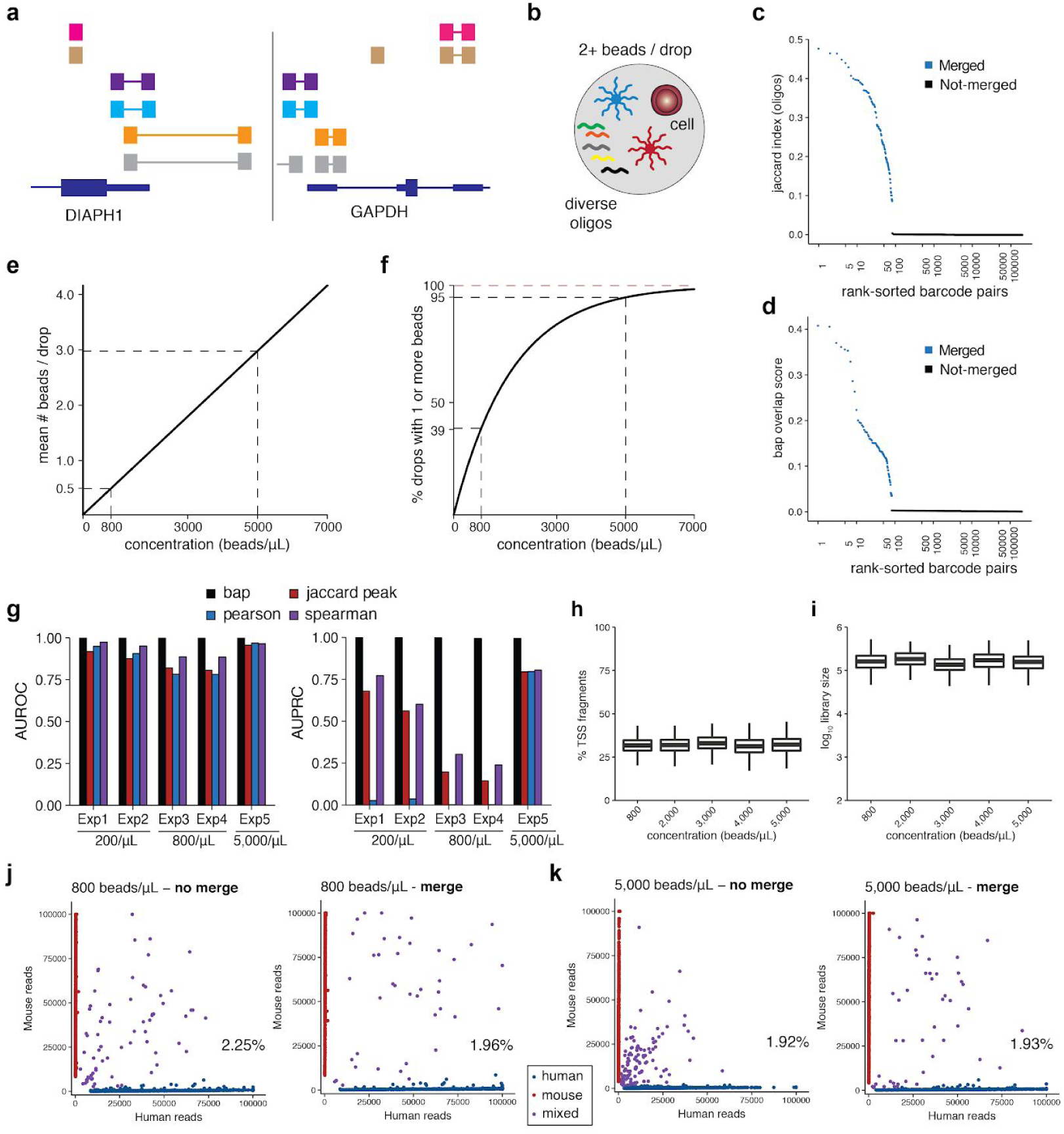
Validation of bead merging computational approach. (**a**) Browser shot of paired-end reads near the *DIAPH1* and *GAPDH* locus. Reads are colored by bead barcode sequence. (**b**) Schematic of verification experiment where a library of random oligonucleotides was encapsulated into droplets together with Tn5 transposed cells and barcoded beads. Schematic shows a droplet containing a library of random oligos, a cell and two beads with different barcode sequences. (**c**) Jaccard index overlap plot for pairs of bead barcodes. For each pair of bead barcodes observed, the Jaccard index was computed over the observed oligonucleotide sequences. (**d**) The bap overlap score computed from the scATAC-seq data (agnostic to oligonucleotides) from the same experiment. In each panel, pairs of bead barcodes nominated for merging are highlighted in blue. Merged pairs were determined by computing a “knee” inflection point. (**e**) The expected number of beads per drop as a function of bead concentration. Inference of this line was determined by a maximum likelihood estimation for a double-truncated Poisson distribution. (**f**) Percent of drops with one or more beads as a function of bead concentration. Values are estimated using the probability density function of a Poisson distribution parameterized by the mean number of beads per drop from (**e**). (**g**) (left panel) Area under the receiver operating curve (AUROC) values for true positive bead merges nominated from the oligonucleotide sequences. Four metrics are compared, including Pearson and Spearman correlation and the Jaccard index of reads in peaks per pair of bead barcodes. The final metric is our novel computational approach, termed bap. Various bead concentrations per experimental condition are shown below the x-axis. (right panel) The same conditions and metrics but showing the area under the precision-recall curve (AUPRC). (**h**) %TSS enrichment scores for the same pool of cells processed at different bead concentrations. Center line, median; box limits, first and third quartiles; whiskers, 1.5x interquartile range. (**i**) Per-cell library complexities across a range of tested bead concentrations, the same as in panel (**h**). Center line, median; box limits, first and third quartiles; whiskers, 1.5x interquartile range. (**j**) Species mixing plots and collision rates (text) for the same experiment (800 beads / μL) with and without bead merging. (**k**) The same plots as in (**j**) but at a bead concentration of 5,000 beads / μL.

**Figure S3.**
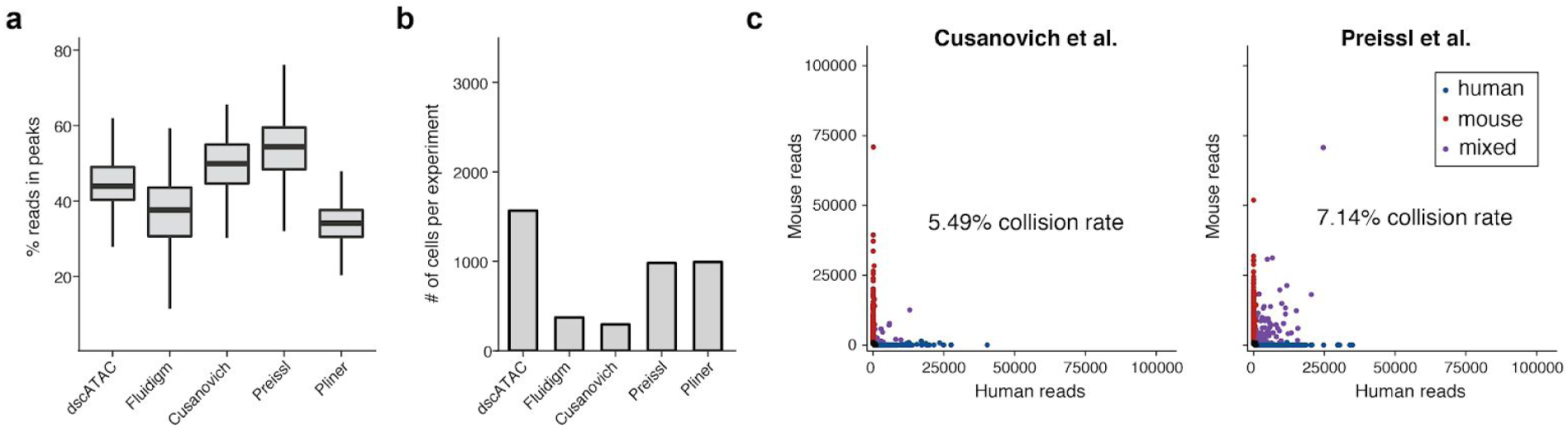
Additional quality controls of dscATAC-seq. (**a**) Fraction of reads in peaks for the comparison in Figure 1d. The chromatin accessibility peak set was obtained from ENCODE DNase-seq data for GM12878 and thus agnostic to the datasets compared here. Center line, median; box limits, first and third quartiles; whiskers, 1.5x interquartile range. (**b**) Number of cells (GM12878 only) compared in panel (**a**) and Figure 1d. (**c**) Species mixing plots and estimated collision rates for existing scATAC-seq methods.

**Figure S4.**
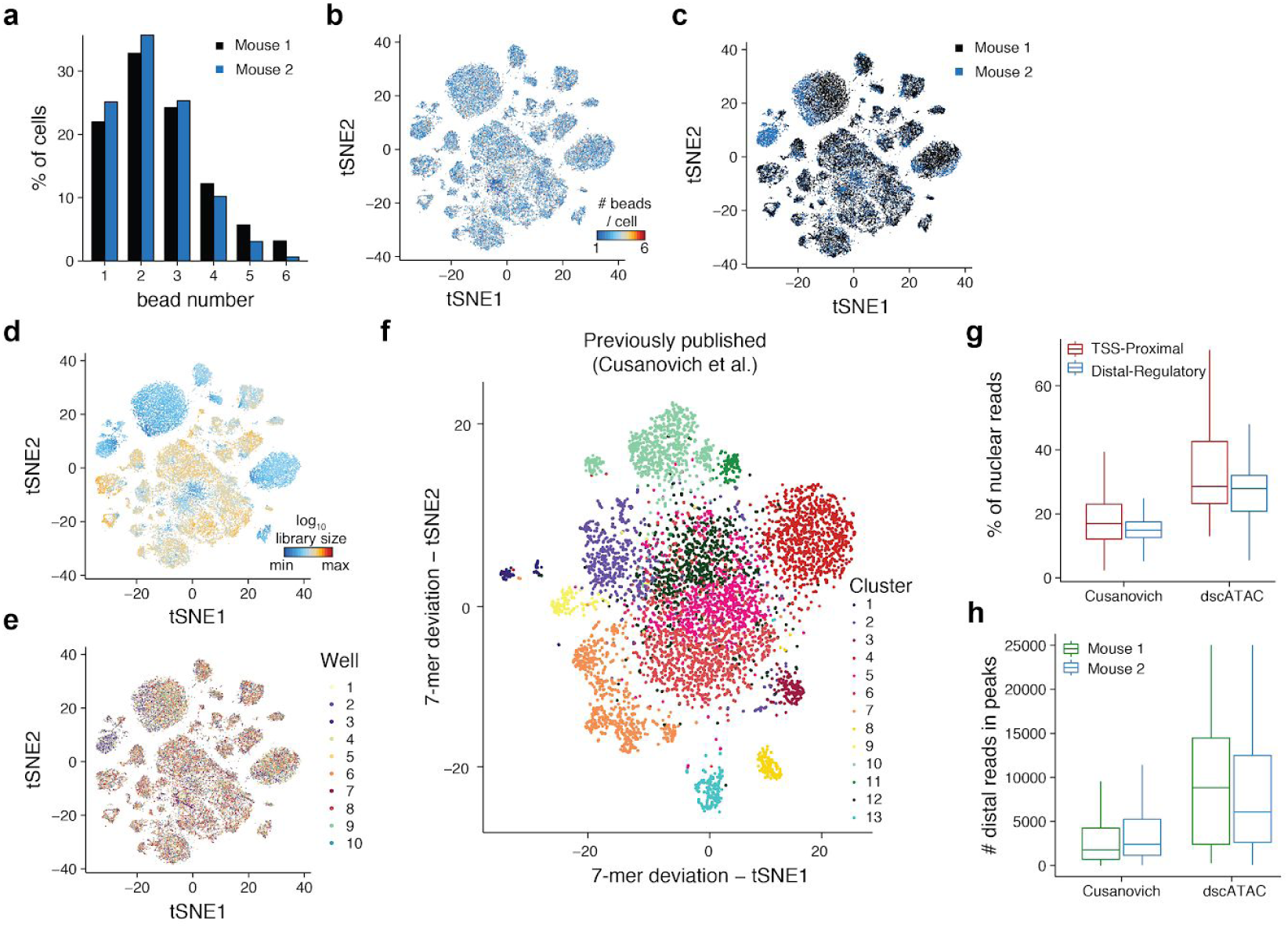
Quality control information for the dscATAC-seq mouse brain dataset and comparison with existing data. (**a**) Distribution of number of beads per cell identified across the two mice (bead input concentration = 5,000 beads / μL) for high-quality cells that pass quality controls. (**b**) Mouse brain cells in the t-SNE from Figure 2a colored by number of bead barcodes detected per cell. The same coordinates are shown for (**c**) mouse donor, (**d**) per-cell log10 library complexity, and (**e**) experimental well. (**f**) t-SNE of previously published sciATAC-seq data for mouse brain (ref. 27) using the same 7-mer method (Louvain, t-SNE; compare to Figure 2a). (**g**) Comparison of the percentage of reads mapping to the nuclear genome (separated into TSS-proximal or distal chromatin accessibility peaks) between whole mouse brain data generated using dscATAC-seq or a recently optimized sciATAC-seq method (ref. 27). Center line, median; box limits, first and third quartiles; whiskers, 1.5x interquartile range. (**h**) Raw total number of reads mapping to distal chromatin accessibility peaks (see blue from panel (**g**) between dscATAC-seq and the sciATAC-seq method described in (**g**). Boxplots summarize thousands of cells for each comparison. Center line, median; box limits, first and third quartiles; whiskers, 1.5x interquartile range.

**Figure S5.**
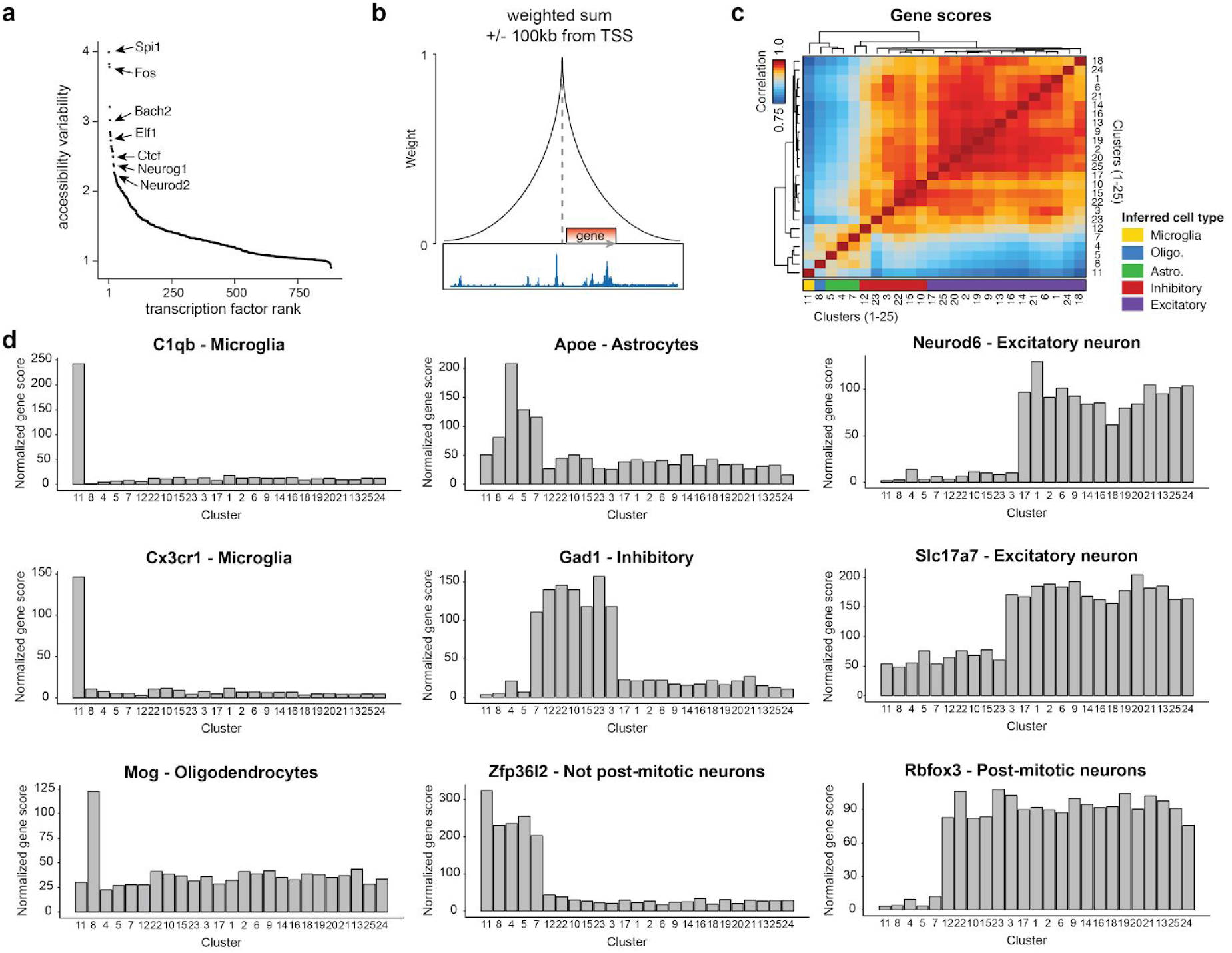
Chromatin accessibility scores for validation of cell clusters from mouse brain. (**a**) Most variable transcription factors identified in the mouse brain dscATAC-seq dataset computed using chromVAR. (**b**) Schematic demonstrating the approach used to define chromatin accessibility scores surrounding gene promoters. (**c**) Hierarchical clustering of chromatin accessibility scores calculated as shown in (**b**) for each cluster derived from the mouse brain dscATAC-seq dataset. (**d**) Representative chromatin accessibility scores for known marker genes defining cell types in the mouse brain, plots are titled by the marker gene and defined cell type.

**Figure S6.**
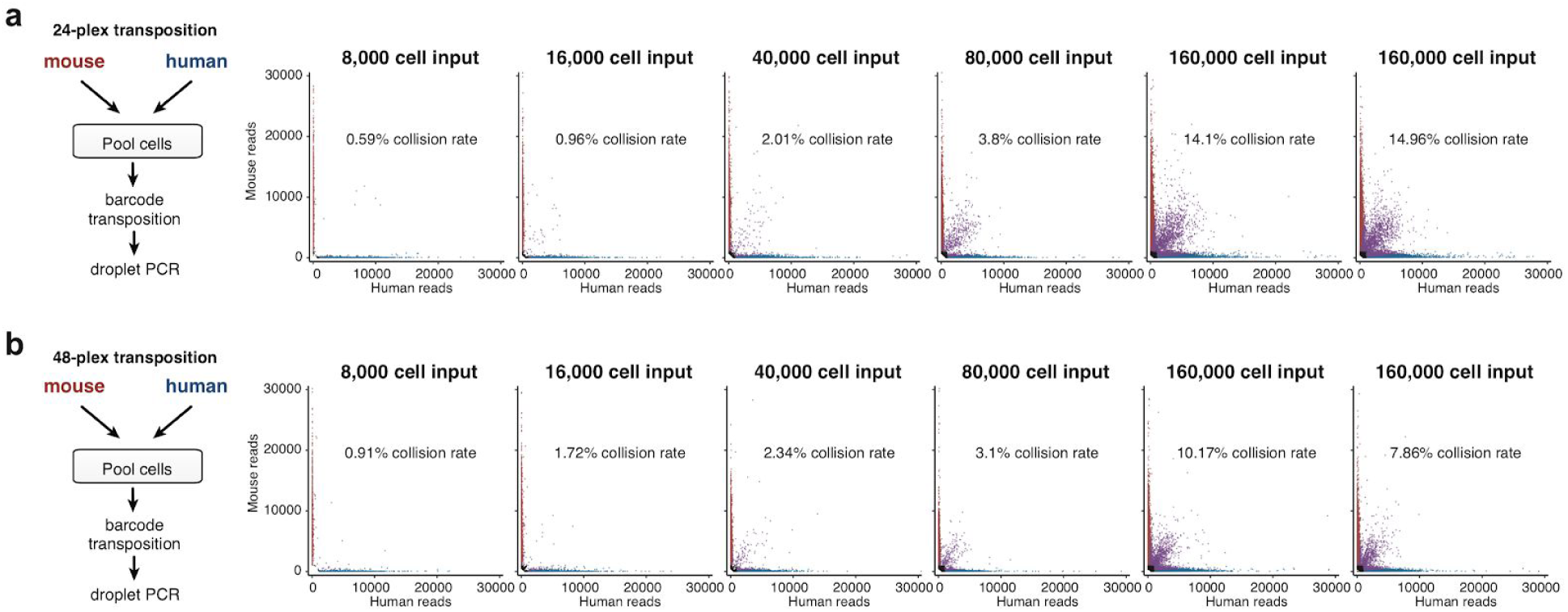
Species mixing analysis of dsciATAC-seq. (**a-b**) Species mixing analysis for human (K562) and mouse (3T3) cell mix generated using (**a**) 24 or (**b**) 48 Tn5 transposase barcodes. For each panel a schematic of the experimental procedure is included (left), and primary results from a cell titration plotting total mouse or human nuclear fragments (right). In these plots points are labeled as either low quality (black), mouse (red), human (blue) or mixed (purple).

**Figure S7.**
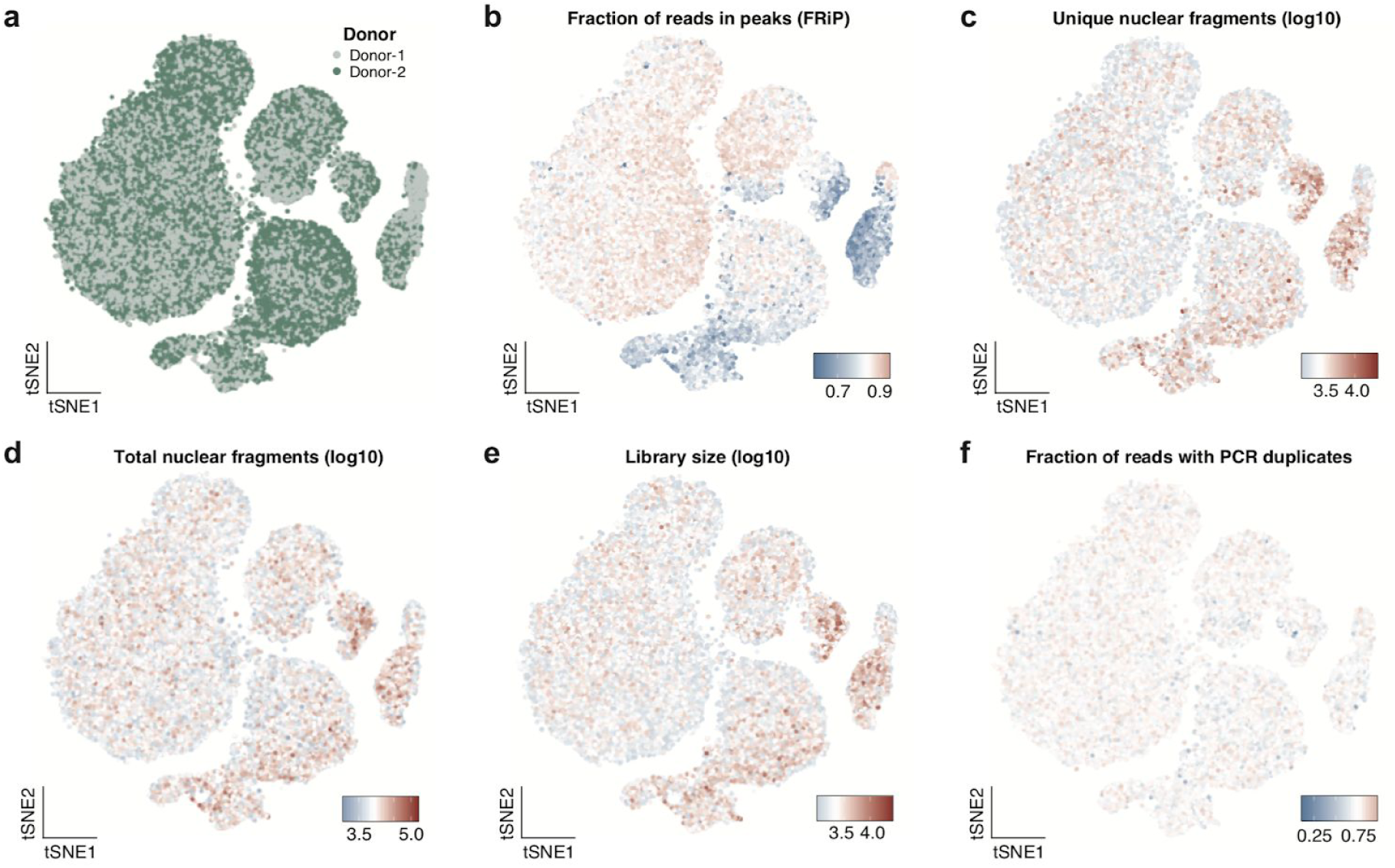
Quality control analysis of human bone marrow dsciATAC-seq data. (**a-f**) Single-cell data derived from BMMCs colored by their (**a**) donor, (**b**) fraction of reads in peaks (FRiP), (**c**) log10 unique nuclear fragments, (**d**) log10 total aligned nuclear fragments, (**e**) log10 library size, and (**f**) fraction of reads with PCR duplicates.

**Figure S8.**
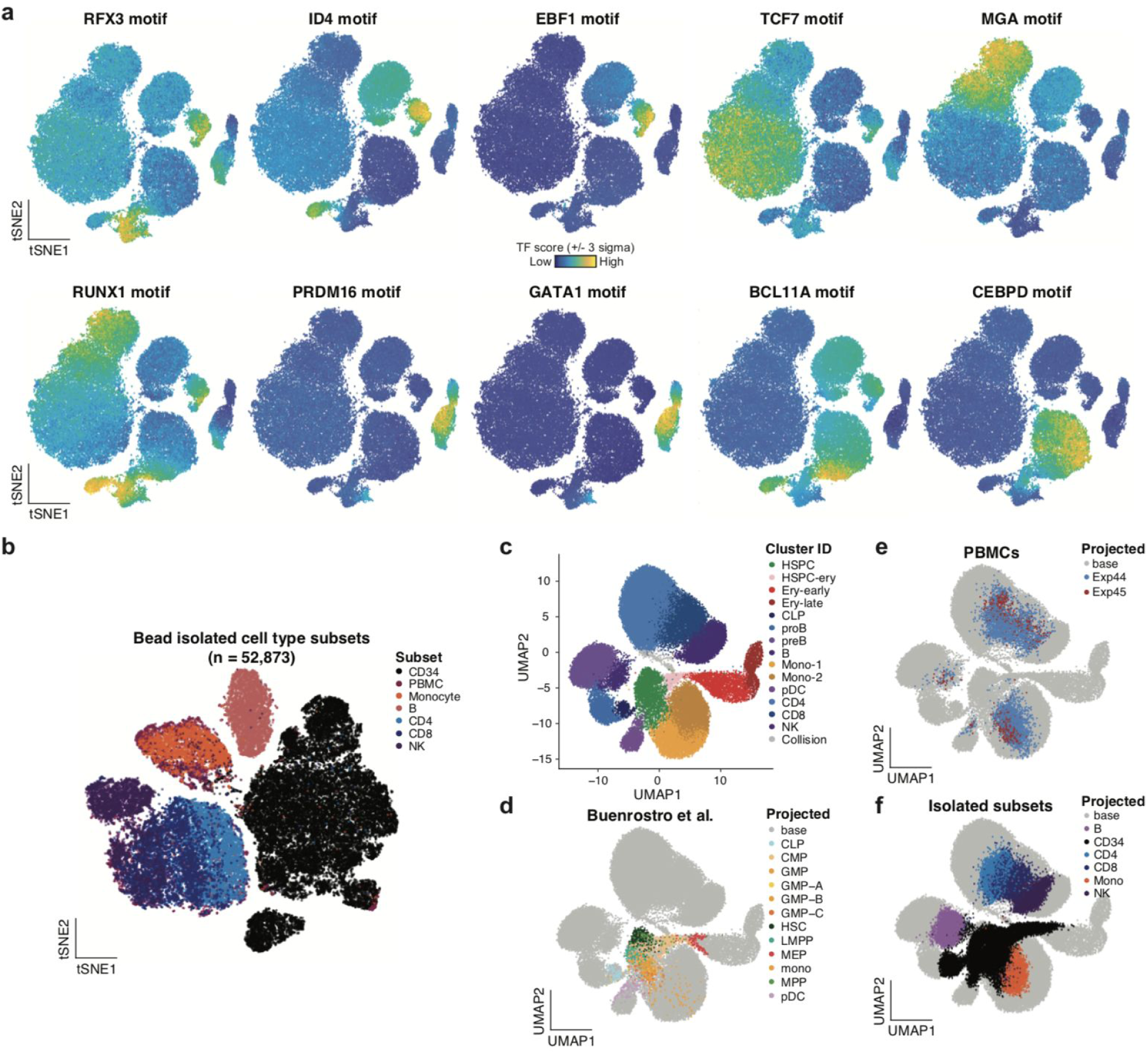
Cell types identified in the human bone marrow dsciATAC-seq data. (**a**) Selected transcription factor deviation motifs shown for resting cells profiled using dsciATAC-seq. (**b**) Embedded cells from isolated subtypes profiled using the standard dscATAC-seq platform. (**c**) UMAP embedding of single-cell data colored by clusters identified (compare to **Fig. 4c**). (**d-f**) Projection of additional single-cell data onto UMAP coordinates of the dsciATAC-seq bone marrow data, projecting (**d**) sorted progenitor subsets (Buenrostro et al., ref. 21), (**e**) peripheral blood mononuclear cells (PBMCs) or (**f**) isolated subsets (shown individually in (**b**)).

**Figure S9.**
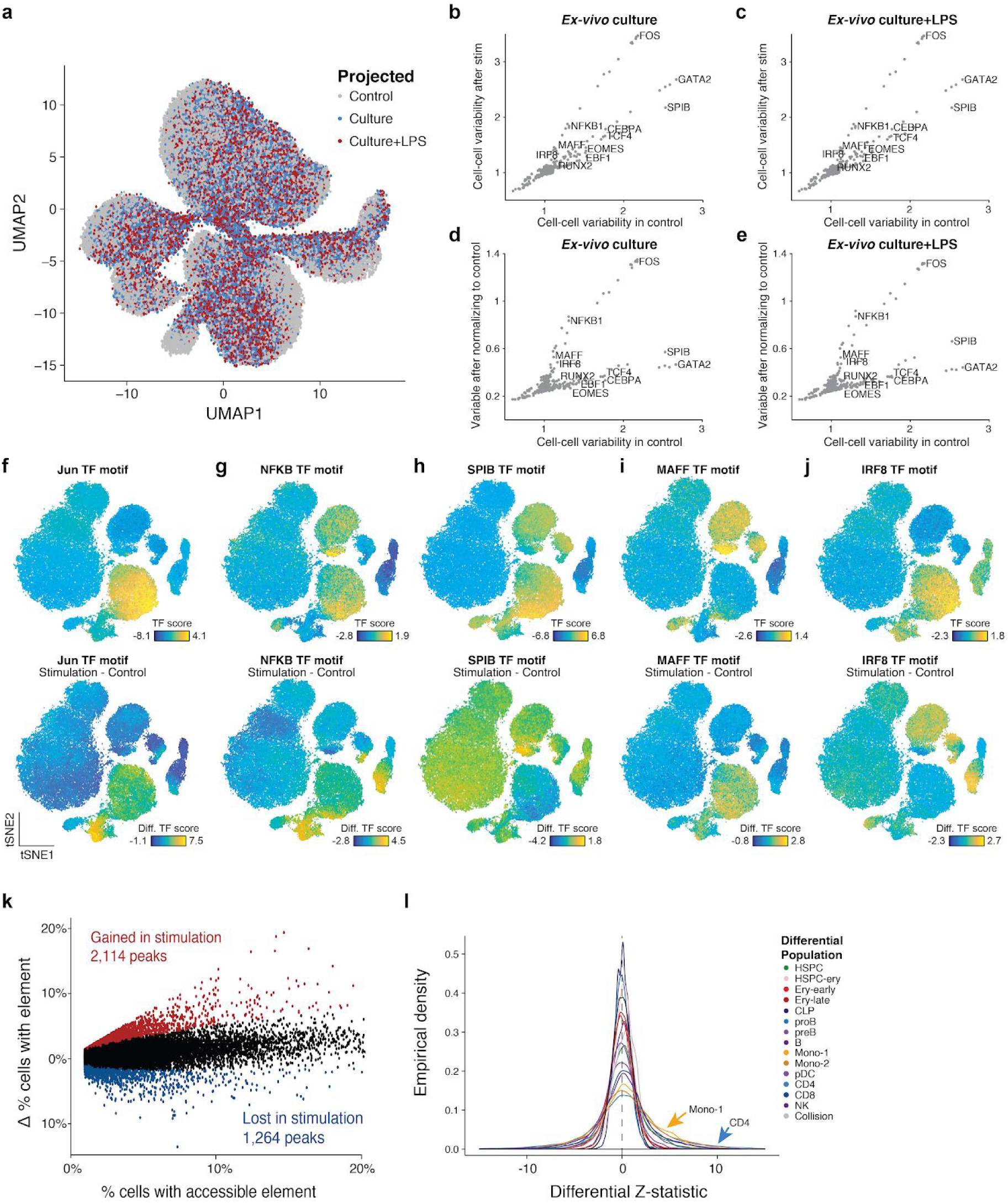
Stimulation of human bone marrow derived cells. (**a**) Stimulated BMMC cells projected onto the UMAP coordinates defined by the non-stimulated control cells. (**b,c**) Cell-cell TF score variability for the stimulation and control cells showing (**b**) *ex vivo* culture and (**c**) *ex vivo* culture and LPS stimulation, only unique TF motifs are highlighted. (**d,e**) Cell-cell TF score variability for the control cells and variability of stimulation after normalizing the control TF variability for (**d**) *ex vivo* culture and (**e**) *ex vivo* culture and LPS stimulation conditions, only unique TF motifs are highlighted. (**f-j**) Depictions of transcription factor deviation scores in resting cells (top) compared to the differential (bottom) after stimulation for selected motifs. (**k**) Sample summary of differential peak analysis for the Mono-1 cluster. Each dot represents a chromatin accessibility peak found in at least 1% of cells. The overall % of cells with element are shown on the x-axis whereas the y-axis depicts the difference in the % of cells with the element accessible (stimulated - resting). Peaks found significantly different at a 1% FDR are colored in red and blue. (**l**) Overall summary statistics per-population from differential peak analysis showing the Z-statistic from the permutation test for differential accessibility. Each colored curve represents the overall Z-statistics for all peaks in the specified cluster.

